# A critical period for prefrontal cortex PV interneuron myelination and maturation

**DOI:** 10.1101/2023.08.15.553393

**Authors:** S. Hijazi, M. Pascual-García, Y. Nabawi, S.A. Kushner

**Affiliations:** Department of Psychiatry, Erasmus MC, Rotterdam, The Netherlands; Department of Pharmacology, University of Oxford, Oxford, UK; Department of Psychiatry, Columbia University, New York, USA

## Abstract

Recent studies have highlighted axonal myelination as a common feature of parvalbumin-positive (PV) interneurons throughout the cerebral cortex. However, the precise function of PV interneuron myelination remains incompletely understood. In this study, we used the cuprizone model of demyelination to investigate how PV interneuron myelination might influence their neuronal physiology. Specifically, we examined whether impairing myelination from postnatal day 21 onwards, during a critical neurodevelopmental period of the prefrontal cortex (PFC), can affect PV interneuron maturation and function. Using whole-cell patch-clamp recordings to examine intrinsic properties of PV interneurons in the PFC, we found that juvenile demyelination induced robust alterations of PV interneuron firing patterns. Specifically, we observed that demyelination caused an impairment in the ability of PV interneurons to sustain high frequency firing associated with a substantial decrease in Kv3-specific currents. We also found a significant impairment in PV interneuron autaptic self-inhibitory transmission, a feature implicated in temporal control of PV interneuron firing during cortical network activity. Following a remyelination period of 5 weeks, PV interneuron properties were only partially recovered and mice showed clear social deficits, suggesting that transient juvenile demyelination leads to long-lasting behavioral impairments. In contrast, adult demyelination had no significant effects on PV interneuron firing properties. Together, our data uncovers a critical period for juvenile myelination as an important factor in PFC PV interneuron development and brain maturation.

## Introduction

Cortical parvalbumin (PV) interneurons, also known as fast-spiking interneurons undergo accelerated developmental maturation, including narrowing of the action potential (AP) waveform and high frequency firing, during a stereotyped postnatal time window extending between postnatal day (P)14 and P28, which depends on the developmentally-regulated expression of a specific complement of voltage-gated ion channels (Ethan M. Goldberg et al. 2011; Daw, Ashby, and Isaac 2007; Micheva et al. 2021b). Another important developmental aspect of PV interneurons is their extensive myelination which emerges during a closely overlapping time window (Micheva et al. 2021b; 2016; J. Stedehouder et al. 2017). While the role of myelination has been widely studied in long-range projection neurons, the impact of myelination on local PV cortical interneurons is still poorly understood. Recent studies have endorsed the classical view that myelination of PV interneurons functions to increase axonal conduction velocity in a manner similar to that of projection neurons (Micheva et al. 2021a), while also being critical for PV interneuron-mediated feedforward inhibition in cortical sensory circuits (Benamer et al. 2020) and modulation of local circuit synchronization (Dubey et al. 2022). However, the specific role of PV interneuron myelination over the time course of neurodevelopment has yet to be elucidated.

Aberrant PV interneuron maturation has been suggested as a contributor to the pathogenesis of multiple neurodevelopmental diseases, and impairments in the development of prefrontal cortex (PFC) circuits has been at the center of many studies investigating neurodevelopmental disorders (J. Stedehouder and Kushner 2017; Leicht et al. 2016; van Os and Kapur 2009; Chini and Hanganu-Opatz 2021; Bicks et al. 2020). Hence, the maturation of PV interneurons in PFC is of particular significance. Notably, PFC development persists long after adolescence (Chini and Hanganu-Opatz 2021; Klune, Jin, and DeNardo 2021) and PFC-dependent functions, such as cognitive flexibility and emotional memory encoding, start emerging later in development (Chini and Hanganu-Opatz 2021; Klune, Jin, and DeNardo 2021; Coley et al. 2021). This delayed period of maturation of PFC circuits underlies an important time window necessary for the establishment of specific neuronal circuits within the PFC^16–18^. Conversely, it also reveals a transient period during which the PFC is vulnerable to various types of stressors and insults (Bitzenhofer et al. 2021).

Here, we investigated how the neurodevelopmental timing of myelination influences PV interneuron maturation in the PFC. Our data reveal that myelination of PV interneurons during a critical period of development is instrumental for PV interneuron maturation and their unique ability to fire at high frequencies. We find that impaired myelination in early adolescence can have long lasting effects on PV interneuron morphology and function and impair social behavior in adulthood.

## Results

### Myelination-dependent critical period for PV interneuron maturation

#### Juvenile demyelination affects PV interneuron properties, maturation and morphology in PFC

In order to investigate the neurodevelopmental relationship between myelination and PV interneuron maturation in the PFC, mice were fed a 0.2% cuprizone diet for 6 weeks starting at P21. Mice were then sacrificed and brain slices prepared for whole-cell patch-clamp recording and filling of PV interneurons (**Figure 1A**). Juvenile cuprizone treatment led to a clear decrease in myelination in adulthood and a complete demyelination of PV interneuron axons in the PFC (**Figure 1B-C**). Next, we fully reconstructed biocytin-filled patched cells and quantified their axonal and dendritic processes (**Figure 1D-E**). We found a significant decrease in total axonal length and axonal branching of PV interneurons from mice that underwent juvenile demyelination (**Figure 1F**). The density of PV interneurons and their total dendritic length were unaffected (**Figure 1G-I**).

**Figure 1.**
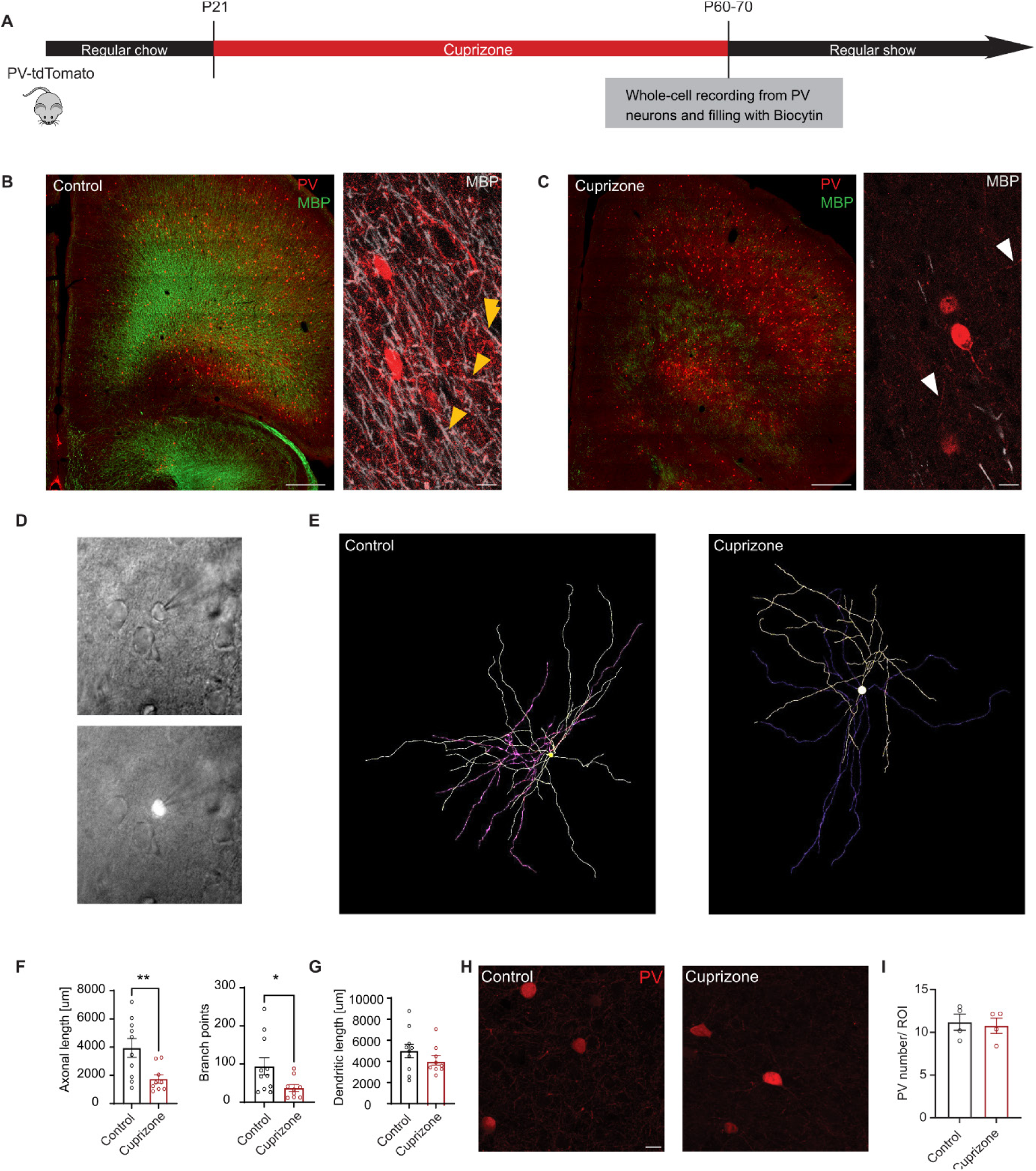
Juvenile demyelination leads to a decrease in axonal complexity of PFC PV interneurons. **A.** Experimental design to induce juvenile demyelination in PV-tdTomato mice **B.** *Left.* Confocal overview image of PFC in PV-tdTomato animal (tdTomato+, red) overlaid with myelin basic protein (MBP, green); scale bar: 300um. *Right*. 40x image showing myelinated PV interneuron axons in control conditions (MBP, white); scale bar: 20um. **C**. *Left.* Confocal overview image of the PFC region immunolabeled for MBP expression showing the loss of myelin following 6 weeks of 0.2% cuprizone-diet. *Right*. 40x image showing no myelination of PV interneuron axons in cuprizone conditions. Yellow arrow heads point to PV interneuron axons. **D**. Live contrast image (upper) of a tdTomato-labeled (lower) PV interneuron in the PFC region with the patch pipette attached. **E**. Example image of a fully reconstructed biocytin-labeled PFC PV interneuron from a control mouse (left) and from cuprizone-treated mouse (right). **F**. *Left.* Total proximal axonal length is decreased in cuprizone-treated PFC PV interneurons compared with control cells (*t-test; n = 10/9 cells per group, **p < 0.01). Right*. The number of branch points in PV interneuron axons from cuprizone-treated mice was significantly decreased compared to control mice (*t-test; n = 10/9 cells per group, *p < 0.05*). **G**. Total dendritic lengths were unaltered by juvenile demyelination (*t-test; n = 10/9 cells per group, p = 0.2856*). **H.** PV interneurons from coronal sections of the PFC of control (left panel) and cuprizone-treated (right panel) mice using anti-PV staining (red). The number of PV-positive cells was quantified in high magnification images. Scale bar: 20 um. **I**. Quantification of the total number of PV interneurons in the PFC region shows no difference between control (black) and cuprizone-treated (red) mice (*t-test; n = 4 mice per group, p = 0.7577*).

Given that demyelination during adolescence led to changes in PV interneuron morphology, we next aimed to characterize whether PV interneuron physiology was correspondingly affected. To that end, we performed current clamp recordings from PFC PV interneurons following juvenile demyelination. Our data revealed alterations in both passive and active membrane properties of PFC PV interneurons in mice that underwent juvenile demyelination, indicative of an immature PV interneuron phenotype (**Figure 2**). Specifically, there was a significant increase in input resistance and sag amplitude (**Figure 2A-F**). The action potential (AP) waveform was also affected (**Figure 2G**), showing a substantial increase in AP width, decay time and afterhyperpolarization (AHP) duration (**Figure 2I-L**). Furthermore, there was a significant decrease in the firing frequency of PV interneurons, along with an impairment in the maximum firing frequency (**Figure 2M-O**). Strikingly, the difference in firing frequency was apparent from 50 Hz onwards (**Figure 2N**), in which 38.4% (20 out of 52) of PV interneurons from mice with juvenile demyelination exhibited a failure to sustain repetitive firing, compared to only 11.6% (5 out of 43) of the cells from the control group (Fisher’s Exact Test, \*\**p*=0.004; **Figure 2P-Q**). Taken together, these data suggest that juvenile demyelination impairs PV interneuron maturation in the PFC.

**Figure 2.**
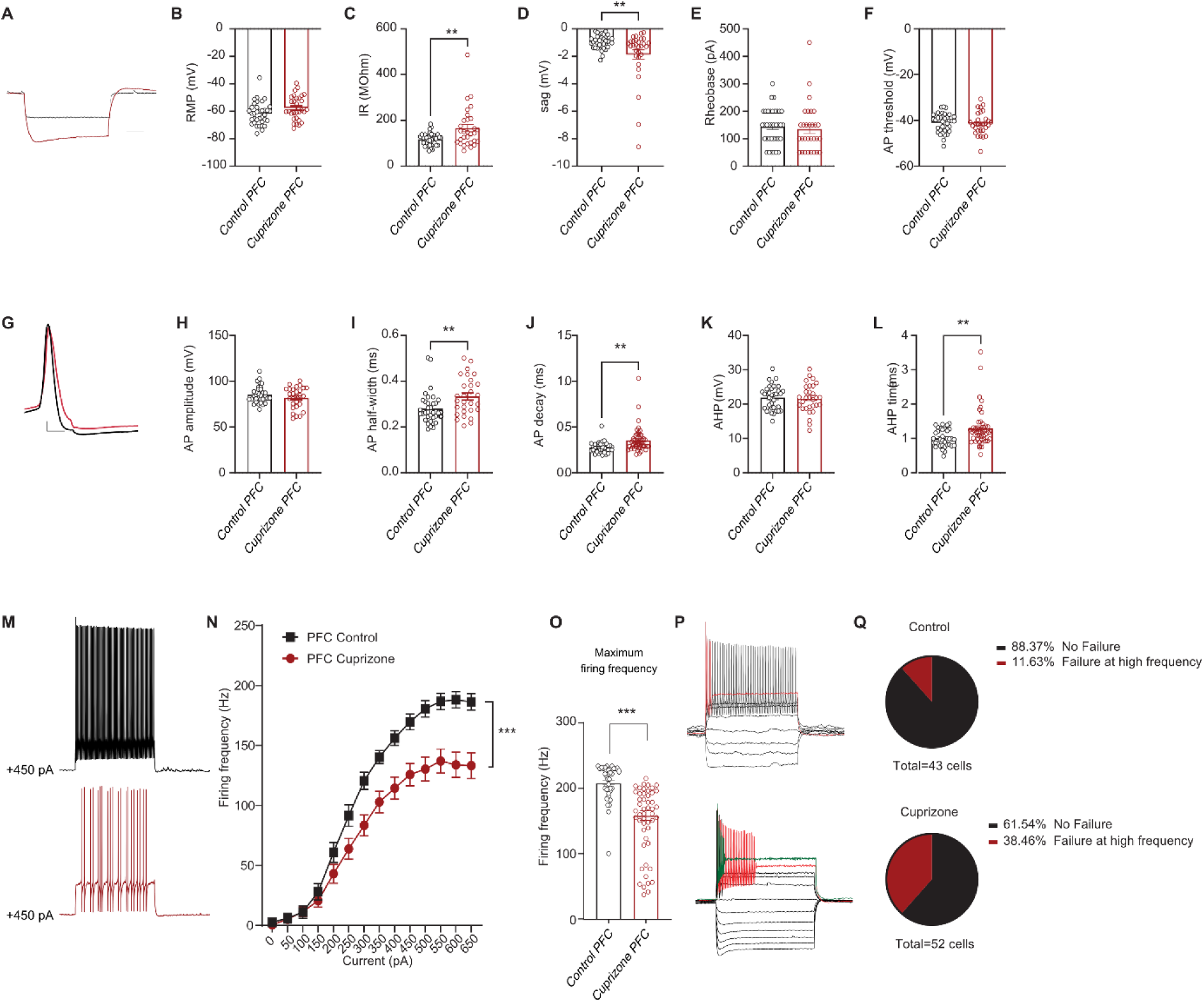
Juvenile demyelination impairs the maturation of PV interneurons in the PFC. **A**. Representative traces of voltage responses following a hyperpolarizing step from control (black) and cuprizone-treated (red) mice illustrating an increased input resistance and sag in mice that underwent juvenile demyelination. Scale bar: 100 ms, 10 mV. **B**-**F**. Summary data showing the averaged (± s.e.m) of the following intrinsic properties: (B) Resting membrane potential (RMP) (*t-test; n = 36/30 cells from 9/8 mice per group, p = 0.0511*), (C) Input resistance (IR) (*t-test; n = 36/30 cells from 9/8 mice per group, **p < 0.01*), (D) Sag (*t-test; n = 36/30 cells from 9/8 mice per group, **p < 0.01*), (E) Rheobase (*t-test; n = 36/30 cells from 9/8 mice per group, p = 0.6853*) and (F) Action potential (AP) threshold (*t-test; n = 36/30 cells from 9/8 mice per group, p = 0.7831*). **G**. A representative trace of a single AP illustrating a wider and slower AP in cuprizone-treated mice (red) compared to control (black). Scale bar: 1 ms. **H**-**L**. Summary data showing the averaged (± s.e.m) of the following AP waveform properties: (H) AP amplitude (*t-test; n = 36/30 cells from 9/8 mice per group, p = 0.1534*), (I) AP half-width (*t-test; n = 36/30 cells from 9/8 mice per group, **p < 0.01*), (J) AP decay (*t-test; n = 36/30 cells from 9/8 mice per group, **p < 0.01*), (K) After-hyperpolarization (AHP) amplitude (*t-test; n = 36/30 cells from 9/8 mice per group, p = 0.7903*) and (L) AHP time (*t-test; n = 36/30 cells from 9/8 mice per group, **p < 0.01*). **M**. Representative traces of voltage responses following +450 pA current injection. **N**. Average action potential (AP) frequency in response to 0-650 pA current steps illustrating a significant decrease in PV interneuron firing frequency in cuprizone-treated mice. (*group x current two-way repeated measures: n = 36/30 cells from 9/8 mice per group: F(13,836)= 4.00, p < 0.001*). **O**. Summary data of the maximum firing frequency per group (*t-test; n = 36/30 cells from 9/8 mice per group, ***p < 0.001*). **P**. Example traces of two different PV interneurons that are unable to maintain high-frequency firing at increased current injection. The upper trace shows a cell that can sustain its firing at low-current but not at high, whereas the lower trace shows a cell that cannot sustain its firing at any current-step. **Q**. Percentage of cells that failed to maintain high frequency firing at > 500 pA in both groups reveals a clear increase in cuprizone-treated mice (*Fisher’s exact test; **p = 0.0045*). Data was obtained from voltage recordings with 500 ms current injections ranging from -300 pA to 650 pA.

#### Shiverer PV interneurons have similar properties to juvenile demyelinated PV interneurons, consistent with the causal requirement for properly timed myelination during PV interneuron developmental maturation

In order to independently confirm the impact of juvenile demyelination on the maturation of PV interneurons in the PFC, we made use of *shiverer* mice, harbouring a germline homozygous deletion of MBP impaired myelination. We performed whole-cell recordings from *shiverer* and wild-type littermate PV interneurons at P60-75 (**Figure 3A-B**). PV interneurons in the PFC of *shiverer* mice exhibited considerable similarity to those of mice that underwent juvenile demyelination (**Figure 3C-N**). Specifically, the sag amplitude, AP half-width, AP decay time and AHP time were all significantly increased compared to WT mice (**Figure 3E,K,L,N**). We found no differences in input resistance, RMP, or rheobase (**Figure 3D,G**). Finally, when we examined the firing properties of PV interneurons, we observed a clear decrease in firing frequency at increased current injections, along with a decrease in the maximum firing frequency in *shiverer* mice (**Figure 3O-S**). This was due to the fact that 37.1% (14 out of 37) of the cells from *shiverer* mice could not sustain high frequency firing compared to 4% (1 out of 25) of their WT littermates (Fisher’s Exact Test, **p=0.004; **Figure 3R,S**).

**Figure 3.**
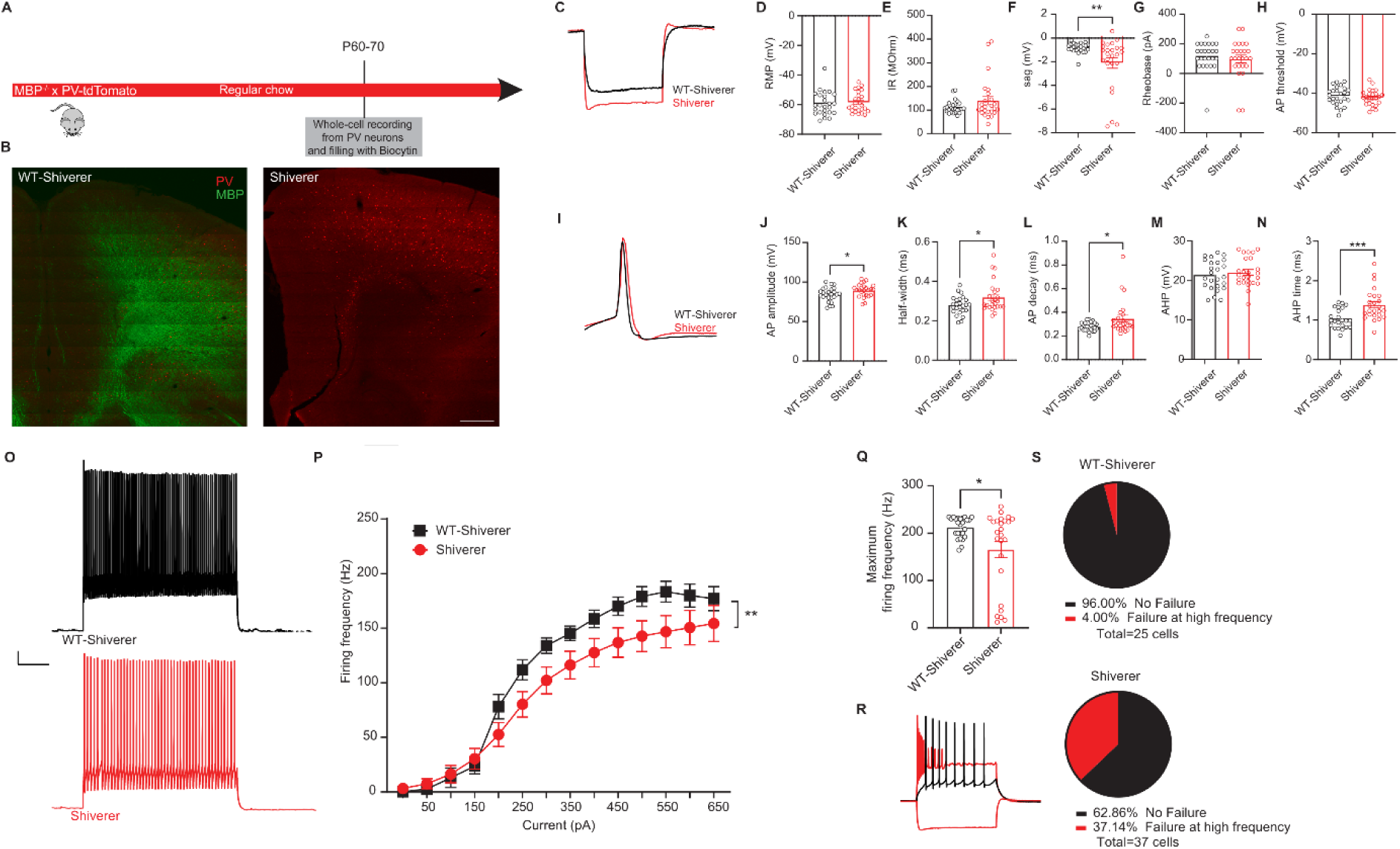
PV interneurons from Shiverer mice show many similarities with the ones from mice that underwent juvenile demyelination. **A.** Experimental design for MBP^-/-^ (Shiverer)-PV-tdTomato mice **B**. *Left.* Confocal overview image of PFC in control PV-tdTomato animal (tdTomato+, red) overlaid with myelin basic protein (MBP, green). *Right.* Confocal overview image immunolabeled for MBP expression showing the absence of myelination in the PFC of Shiverer mice. Scale bar: 300 um. **C**. Representative traces of voltage responses following a hyperpolarizing step from control (black) and Shiverer (light red) mice illustrating an increased sag in mice lacking MBP from birth. **D**-**H**. Summary data showing the averaged (± s.e.m) of the following intrinsic properties: (D) Resting membrane potential (RMP) (*t-test; n = 25/25 cells from 9 mice per group, p = 0.6752*), (E) Input resistance (IR) (*t-test; n = 25/25 cells from 9 mice per group, p = 0.1407*), (F) Sag (*t-test; n = 25/25 cells from 9 mice per group, **p = 0.0072*), (G) Rheobase (*t-test; n = 25/25 cells from 9 mice per group, p = 0.5406*) and (H) Action potential (AP) threshold (*t-test; n = 25/25 cells from 9 mice per group, p = 0.4461*). **I**. A representative trace of a single AP illustrating a wider and slower AP in Shiverer mice (light red) compared to control (black). **J-N**. Summary data showing the averaged (± s.e.m) of the following AP waveform properties: (J) AP amplitude (*t-test; n = 25/25 cells from 9 mice per group, *p = 0.0405*), (K) AP half-width (*t-test; n = 25/25 cells from 9 mice per group, *p = 0.0278*), (L) AP decay (*t-test; n = 25/25 cells from 9 mice per group, *p = 0.0258*), (M) After-hyperpolarization (AHP) amplitude (*t-test; n = 25/25 cells from 9 mice per group, p = 0.5188*) and (N) AHP time (*t-test; n = 25/25 cells from 9 mice per group, ***p = 0.0007*). **O**. Representative traces of voltage responses following +450 pA current injection. **P**. Average action potential (AP) frequency in response to 0-650 pA current steps illustrating a significant decrease in PV interneuron firing frequency in Shiverer mice (*group x current two-way repeated measures: n = 25/25 cells from 9 mice per group: F(13,624)= 2.303, **p < 0.0056*). **Q**. Summary data of the maximum firing frequency per group (*t-test; n = 25/25 cells from 9 mice per group, *p = 0.0109)*. **R**. Example trace of a PV interneuron that is unable to maintain high-frequency firing at increased current injection. **S**. Percentage of cells that failed to maintain high frequency firing at > 500 pA in both groups reveals a clear increase in Shiverer mice (*Fisher’s exact test; **p = 0.0041*).

#### Adult demyelination has no significant effect on PV interneuron’s firing abilities

Are the observed firing impairments at high frequencies due to the loss of myelin during a critical period of development or solely caused by the demyelination of PV interneurons axons? To address this, we recorded from PV interneurons from mice that underwent adult demyelination, in which cuprizone treatment was initiated at P60 instead of P21 (**Figure 4A-C**). PV interneurons from mice with adult demyelination showed a decreased excitability of PV interneurons following adult demyelination with no alterations in firing properties (**Figure 4D, G**). Importantly, input resistance and sag amplitude remained unchanged following adult demyelination (**Figure 4E, F**). Furthermore, the AP waveform and firing frequency of PV interneurons was also unaffected by adult demyelination, in contrast to what was observed following juvenile demyelination and in *shiverer* mice (**Figure 4I-N, Supplementary Figure 1**). Finally, the maximum firing frequency of PV interneurons following adult demyelination also remained intact (**Figure 4P-Q**). Specifically, only 10.0% (2 out of 20) of cells from mice with adult demyelination showed a failure to sustain high firing frequency at high current injections, which was similar to control mice (4.0%, 1 out of 25) (Fisher’s Exact Test, p=0.577; **Figure 4R-S**). Together, our data indicates that adult demyelination has no effect on the AP waveform and the sustained firing properties at high frequencies of PV interneurons, further validating that the aberrant firing observed in the juvenile demyelination group is due to the loss of myelin at a critical age of PFC development.

**Figure 4.**
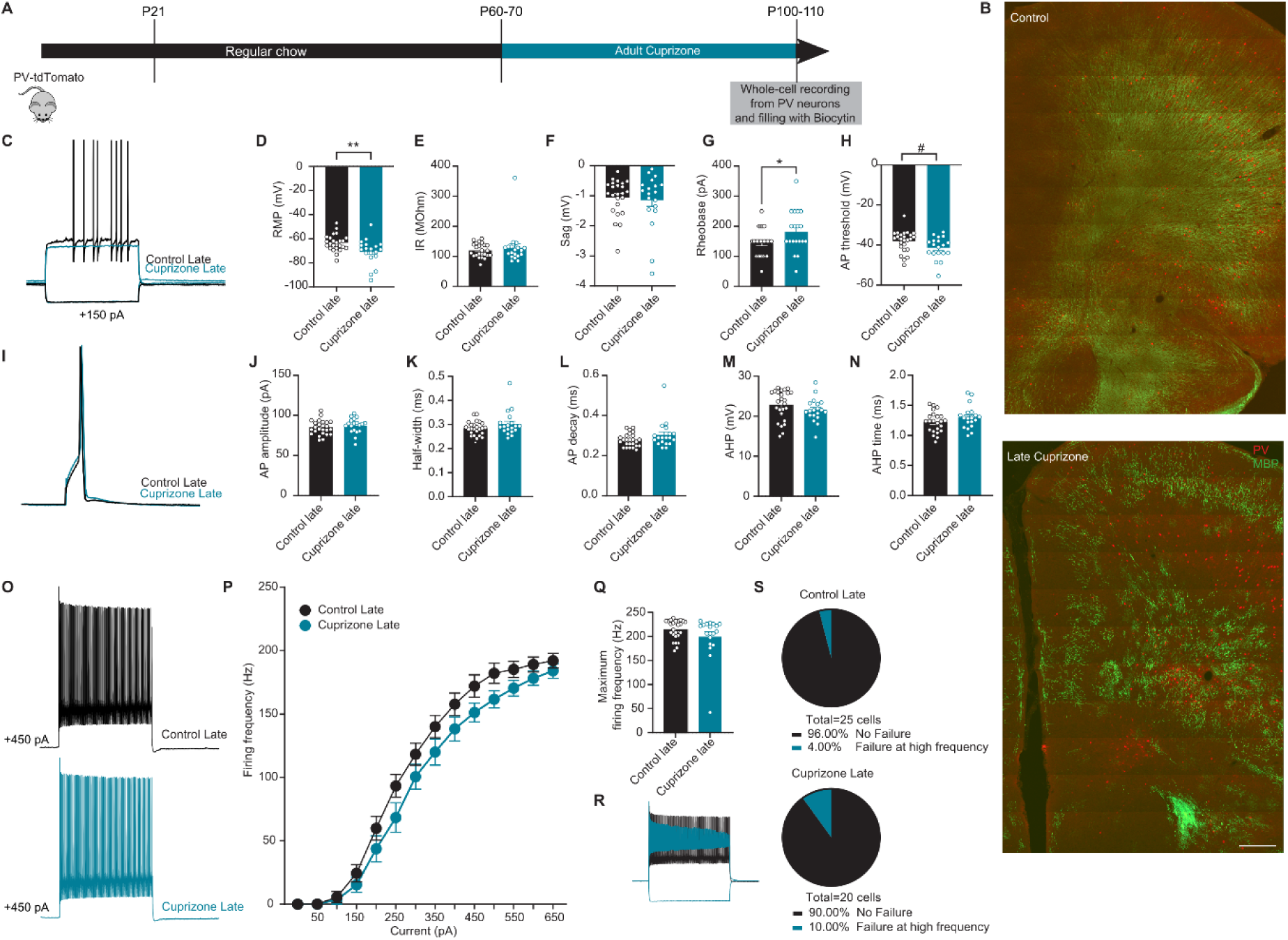
Adult demyelination has no effect on PV interneuron’s firing rate or failures in autaptic responses. **A.** Experimental design for adult demyelination in PV-tdTomato mice. **B**. *Upper panel.* Confocal overview image of PFC in PV-tdTomato animal (tdTomato+, red) overlaid with myelin basic protein (MBP, green). *Lower panel.* Confocal overview image immunolabeled for MBP expression showing the effect of adult demyelination on the PFC region following 6 weeks of 0.2 % cuprizone diet. Scale bar: 300 um. **C**. Representative traces of voltage responses following a depolarizing step from control (Control Late, black) and adult demyelination (Cuprizone Late, blue) mice illustrating an increase in Rheobase in mice following late cuprizone treatment. **D**-**H**. Summary data showing the averaged (± s.e.m) of the following intrinsic properties: (D) Resting membrane potential (RMP) (*t-test; n= 24,19 cells from 8-6 mice per group, **p = 0.0086*), (E) Input resistance (IR) (*t-test; n= 24,19 cells from 8-6 mice per group, p = 0.5066*), (F) Sag (*t-test; n= 24,19 cells from 8-6 mice per group, p = 0.6607*), (G) Rheobase (*t-test; n= 24,19 cells from 8-6 mice per group, *p = 0.0336*) and (H) Action potential (AP) threshold (*t-test; n= 24,19 cells from 8-6 mice per group, #p = 0.0511*). **I**. A representative trace of a single AP illustrating no difference in AP waveform between late cuprizone (blue) compared to control (black). **J**-**N**. Summary data showing the averaged (± s.e.m) of the following AP waveform properties: (J) AP amplitude (*t-test; n= 24,19 cells from 8-6 mice per group, p = 0.3837*), (K) AP half-width (*t-test; n= 24,19 cells from 8-6 mice per group, p = 0.1139*), (L) AP decay (*t-test; n= 24,19 cells from 8-6 mice per group, p = 0.1004*), (M) After-hyperpolarization (AHP) amplitude (*t-test; n= 24,19 cells from 8-6 mice per group, p = 0.2520*) and (N) AHP time (*t-test; n= 24,19 cells from 8-6 mice per group, p = 0.1543*). **O**. Representative traces of voltage responses following +450 pA current injection. **P**. Average action potential (AP) frequency in response to 0-650 pA current steps showing no difference in PV interneuron firing frequency in late cuprizone (blue) mice compared to control (black) mice. (*group x current two-way repeated measures: n = 24/19 cells from 8-6 mice per group: F (13,455) = 1.396, p = 0.1569*). **Q**. Summary data of the maximum firing frequency per group (*t-test; n= 24,19 cells from 8-6 mice per group, p = 0.1581*). **R**. Example trace of a PV interneuron that is unable to maintain high-frequency firing at increased current injection. **S**. Percentage of cells that failed to maintain high frequency firing at > 500 pA was very low after adult demyelination (*Fisher’s exact test; p = 0.5772)*.

### What are the mechanisms underlying impaired high-frequency firing of PV interneurons following juvenile demyelination?

#### PV interneurons show a significant decrease in Kv3 currents and Kv3 expression following juvenile demyelination

Next, we sought to unravel the mechanism by which juvenile demyelination could be impairing PV interneurons firing at high frequency. The ability of PV interneurons to sustain high frequency firing results from their very fast AP properties endowed by a prominent expression of Kv3 channels (Rudy and McBain 2001). Previous studies have identified an upregulation of K+ channel subunits of the Kv3 subfamily during the second and third postnatal week that drives these specific features of fast-spiking PV interneurons (Ethan M. Goldberg et al. 2011; Miyamae et al. 2017). Therefore, we aimed to examine whether the immature phenotype displayed by PV interneurons following juvenile demyelination might be explained by changes in K+ currents in these cells. To that end, we performed voltage-clamp recordings from PFC PV interneurons following juvenile demyelination (**Figure 5**). The amplitude of the K+ currents from +10 mV to +60 mV was significantly lower in PV interneurons following juvenile demyelination (**Figure 5A-B**). To verify that these changes in K+ currents were at least partly due to changes in Kv3 channel activity, we performed the same voltage-clamp recordings in the presence of TEA (1 mM), which results in a relatively selective block of Kv3 potassium channel (Johnston, Forsythe, and Kopp-Scheinpflug 2010) (**Figure 5C**). The data confirmed a decrease in TEA-sensitive currents in the cuprizone group, with a significant two-way interaction of genotype and voltage step (**Figure 5D**). Next, we used immunolabeling against Kv3.1b channels to quantify Kv3 channel expression in PV interneurons from the cuprizone group (**Figure 5E**). We observed a significant reduction of Kv3 immunofluorescence along PV interneurons of the PFC in mice that underwent juvenile demyelination compared to controls, while PV expression and cell density were unaltered (**Figure 5F-J**, **Figure 1H,I**). Together, our results uncover that juvenile demyelination leads to a decrease in Kv3 channel expression and activity within PV interneurons of the PFC.

**Figure 5.**
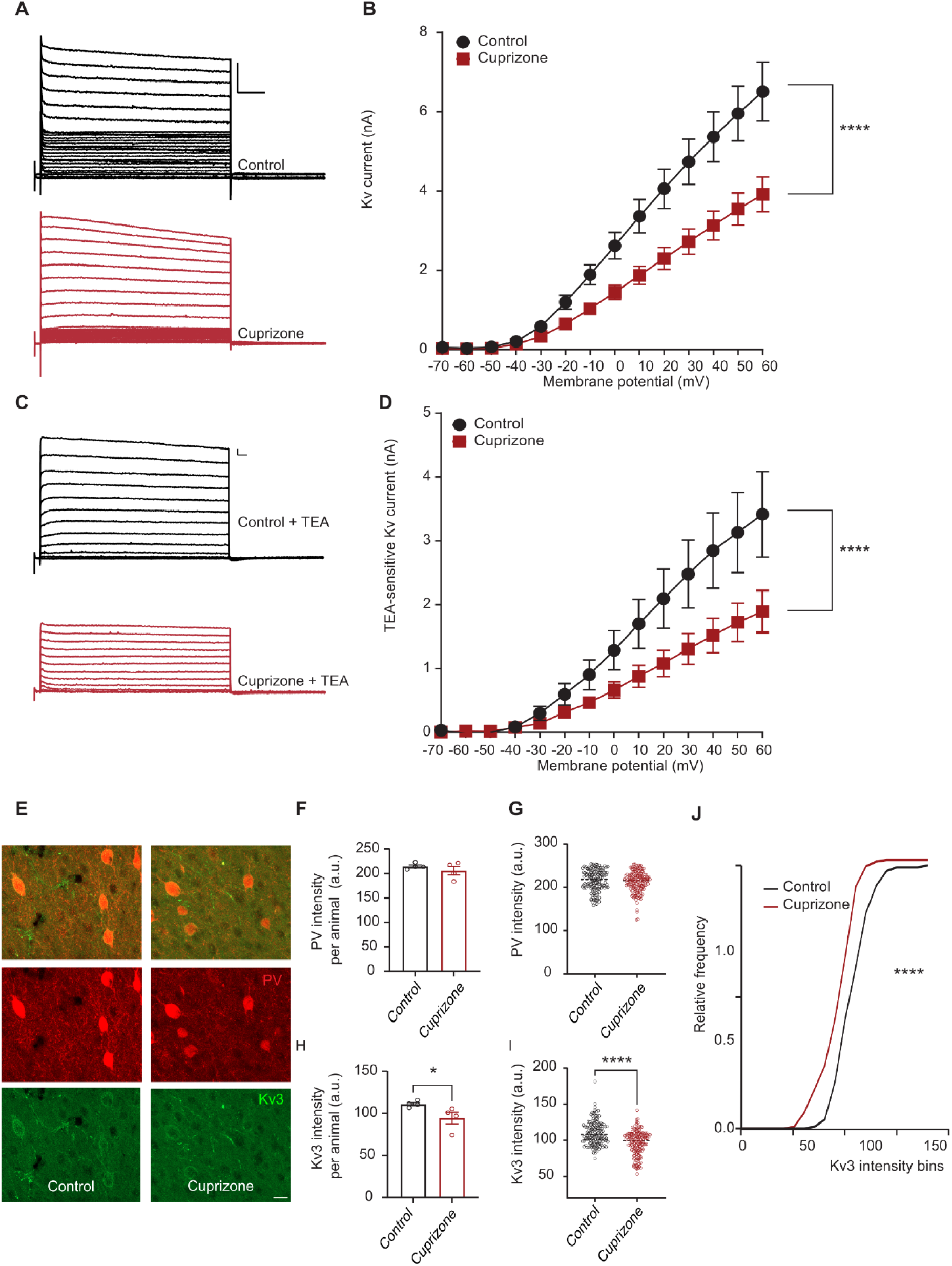
Juvenile demyelination leads to a substantial decrease in potassium currents in PV interneurons of the PFC. **A.** Representative traces of potassium currents evoked with 10 mV potential steps from -70mV to +60 mV in PFC PV interneurons from control (black) and cuprizone (red) mice. Scale: 500 pA, 100 ms. **B**. I–V curves showing a significant decrease in Kv amplitude in PV interneurons from mice that underwent juvenile demyelination. (*group x voltage two-way repeated measures: n = 12,18 cells from 5 mice per group: F(13,364)= 10.45, p < 0.0001).* **C**. Representative traces of TEA-sensitive potassium currents (1mM). **D**. TEA-sensitive currents were specifically decreased in PV interneurons from mice that received cuprizone treatment compared to control mice (*group x voltage two-way repeated measures: n = 10,14 cells from 5 mice per group: F(13,286)= 5.082, p < 0.0001*). **E**. Representative images of double immunostaining of PV interneurons (red) and Kv3.1b (green) in the PFC in control and cuprizone mice. **F**-**G**. PV intensity was not altered following juvenile demyelination (*t-test of (F); n = 4 mice per group, p = 0.3888; t-test of (G); n = 145/170 cells from 4 mice per group, p = 0.3067*). **H**-**J**. Kv3.1b intensity was significantly decreased in PV interneurons from cuprizone mice (red) compared to control mice (black) (*t-test of (H); n = 4 mice per group, p = 0.0286; t-test of (I); n = 145/170 cells from 4 mice per group, p < 0.0001, Kolmogorov-Smirnov test of (J); p < 0.0001*).

#### Modulation of Kv3 channels results in a partial rescue of PV interneuron properties in mice that underwent juvenile demyelination

In order to test whether decreased Kv3 activity contributes to the observed phenotypes reported in PV interneurons after juvenile demyelination, we used AUT00201, a positive modulator of Kv3 potassium currents (Rosato-Siri et al. 2015), to evaluate whether PV interneuron properties are causally related to the observed reduction of Kv3 channel expression following juvenile demyelination. To that end, we measured intrinsic properties of PV interneurons at baseline and following 5 min of bath application of AUT00201 (1uM) (**Figure 6**). We found that AUT00201 decreased the rheobase and rescued AP half-width of PV interneurons from mice with juvenile demyelination (**Figure 6A,B**). Intriguingly, AUT00201 decreased the firing threshold of PV interneurons in both groups, while the remaining properties of PV interneurons were unaltered (**Supplementary Figure 2**).

**Figure 6.**
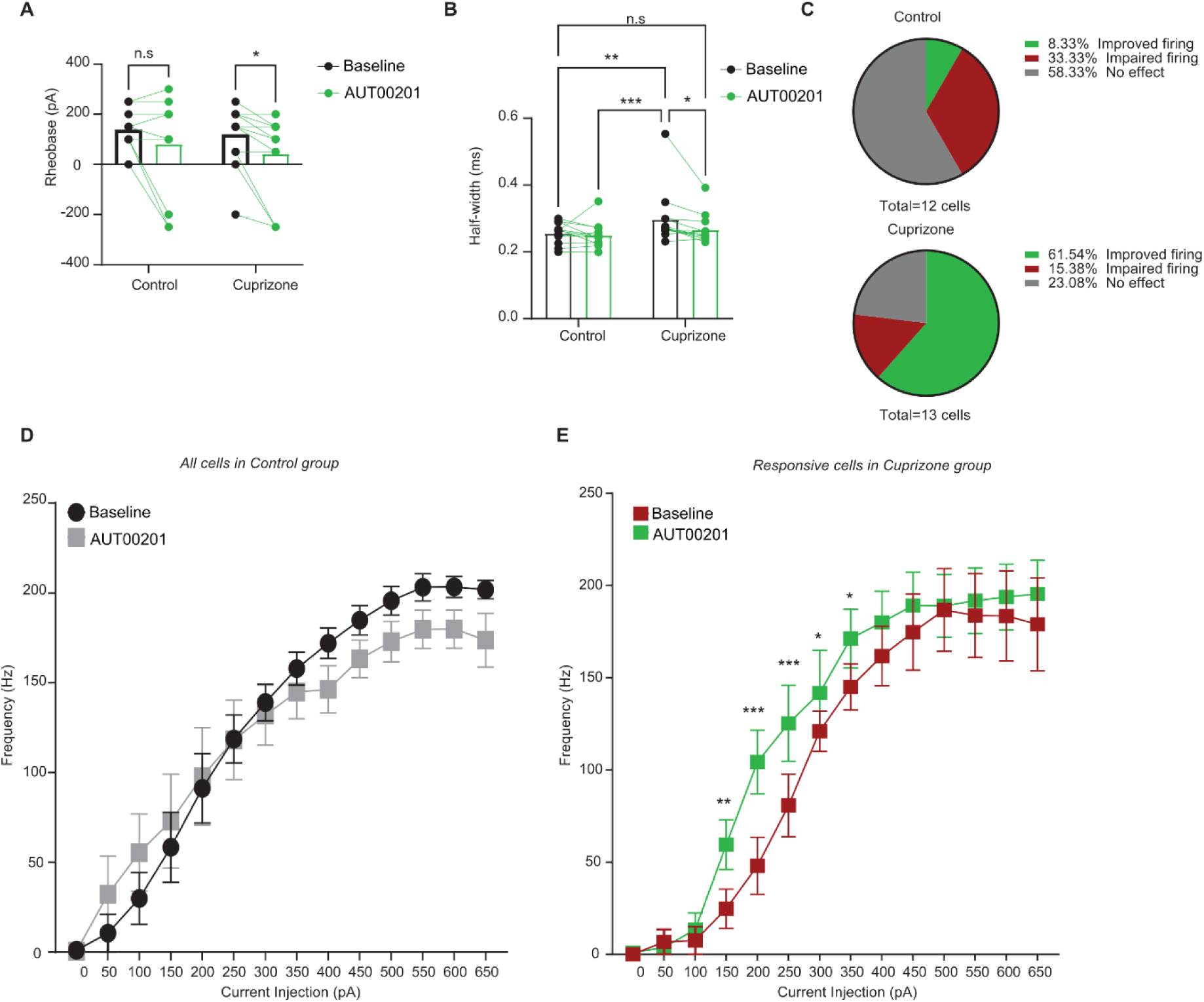
AUT00201 (1uM) can rescue AP width in cuprizone mice. **A**. Bath application of AUT00201 1um decreased the rheobase of PV interneurons from cuprizone mice but not control mice (*group x AUT00201 two-way repeated measures: n = 12,13 cells from 4 mice per group: F(1,12)= 0.1816, p = 0.6775; AUT00201 effect: F(1,12)= 7.122, *p = 0.0205. Post-hoc LSD test: *p < 0.05, n.s. p = 0.0957*). **B**. AUT00201 restores AP half-width in PV interneurons from cuprizone mice, while having no effect on cells from control mice (*group x AUT00201 two-way repeated measures: n = 12,13 cells from 4 mice per group: F(1,12)= 2.774, p = 0.1217; AUT00201 effect: F(1,12)= 5.685, *p = 0.0345. Post-hoc LSD test: *p < 0.05, **p < 0.01, ***p < 0.001, n.s. p = 0.3148*). **C**. Pie charts reflecting the effect of AUT00201 on PV interneurons firing frequency in control (upper) and cuprizone (lower panel) mice. **D**. Average action potential (AP) frequency in response to 0-650 pA current steps suggesting an impairment at high frequencies in control cells after bath application of AUT00201 (1uM) (grey) (*group x current two-way repeated measures: n = 13 cells from 4 mice: p = 0.1369)*. **E**. In the responsive cells from cuprizone mice (red) a significant increase in PV interneuron firing frequency at lower current steps (green) was observed (*group x current two-way repeated measures: n = 8 cells from 4 mice: p = 0.05287; AUT00201 effect: F(1,60)= 5.050, *p=0.0457. Post-hoc LSD test: *p < 0.05, **p < 0.01, ***p < 0.001*).

As AUT00201 rescued AP half-width in mice with juvenile demyelination, we examined whether the I-V relationship of PV interneurons could be rescued by bath application of AUT00201 (1uM). However, when we looked at the firing frequency curves, we did not see any overall significant improvement (**Supplementary Figure 2F**). After a close examination of the effect on AUT00201 on individual cell firing patterns, we found that it increased firing in 61.5 % (8 out of 13) of PV interneurons from juvenile demyelination mice while having no effect in 23.0% of cells (3 out of 13) and impairing (decreasing) the firing in 15.4% of the cells (2 out of 13) (**Figure 6C, Supplementary Figure 3**). Conversely, in control cells, AUT00201 (1uM) increased PV interneuron firing in only 8.3% of the cells (1 out of 12), whereas it had no effect on the firing properties in 58.3% of the cells (7 out of 12). AUT00201 even induced an impairment in firing at high current injections in 33.3% of cells from control mice (4 out of 12) (Fisher’s Exact Test, *p=0.024; **Figure 6C-D**). Analysis of only the positively affected cells in the juvenile demyelination group revealed a significant increase in the firing frequency of PV interneurons after AUT00201 application (**Figure 6E**). Interestingly, bath application of the Kv3 modulator AUT00201 (1uM) had no effect on PV properties in the adult demyelination group (**Supplementary Figure 4**). Taken together, these data imply that a change in Kv3 activity might account for some of the impairment observed after juvenile demyelination in PV interneurons inability to fire at high frequencies.

#### Juvenile demyelination induces a loss of functional autapses and impairs autaptic plasticity

Recent studies have supported a role for autapses, synaptic inputs from individual PV interneurons onto themselves, in regulating their fast-spiking properties and modulation of network oscillations (Deleuze et al. 2019; Deleuze, Pazienti, and Bacci 2014; Szegedi et al. 2020). Accordingly, in the next set of experiments, we set out to explore whether juvenile demyelination altered autaptic transmission, possibly underlying the observed impairments in sustained repetitive firing. To that end, we recorded inhibitory autaptic postsynaptic responses in PFC PV interneurons at P60-75 (**Figure 7A-D**). Significantly fewer PV interneurons from mice with juvenile demyelination exhibited an autaptic response (53.8%; 28 out of 52) compared to the control group (75.9%; 44 out of 58) (Fisher’s Exact Test, *\*p=*0.017; **Figure 7B**). However, among PV interneurons with an autaptic response, autaptic transmission appeared normal (**Figure 7C-D**).

**Figure 7.**
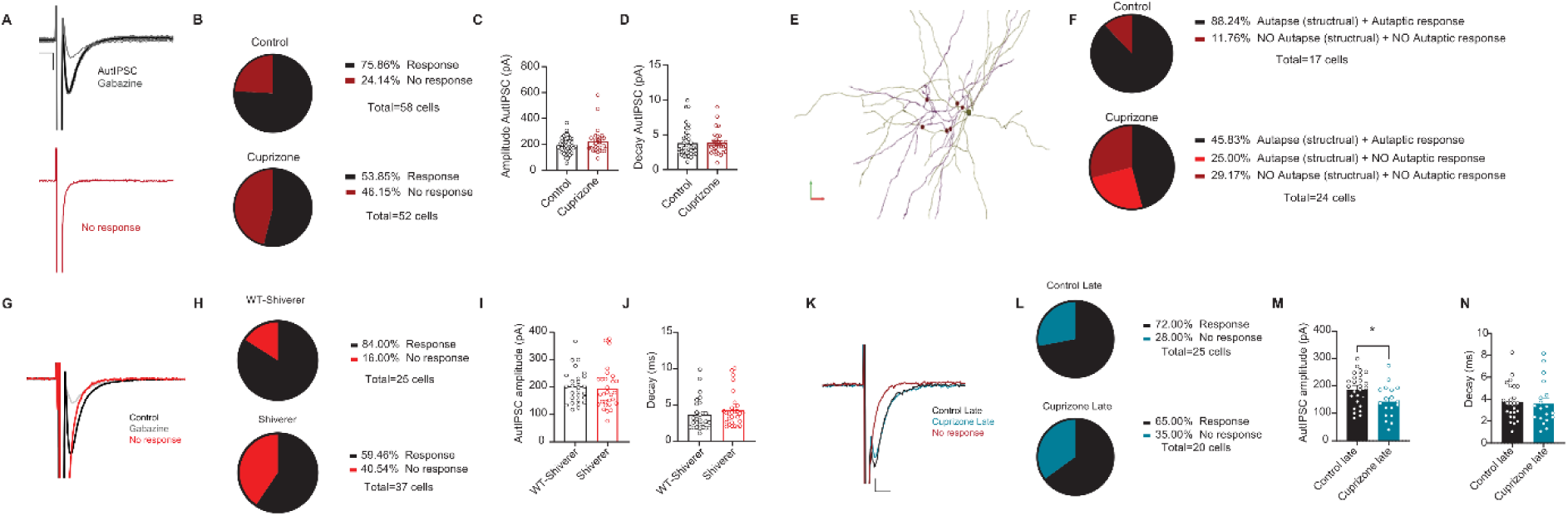
Juvenile demyelination induces a loss of functional autapses. **A**. *Upper panel*. Example traces (average of 8 sweeps) of unitary autaptic inhibitory postsynaptic currents triggered by voltage steps from holding potential of-70mV to 0 mV in PV interneuron (fast inward Na+ currents were partially removed), which can be blocked by gabazine (grey). Note the expected fixed latency for all single-trial responses. *Lower panel*. Example trace showing a failure in autaptic response. scale: 100 pA, 5ms. **B**. Cuprizone-treated mice show a significant decrease in the percentage of cells showing an autaptic response (*Fisher’s exact test; *p = 0.0173*). **C-D**. There was no difference between groups in the amplitude (**C**) or decay (**D**) of the autaptic post-synaptic currents in cells that had autapses (*t-test; n = 40/28 cells per group, p = 0.1246 (C) and p = 0.9350 (D)*). **E**. An example image of a PFC PV interneuron fully reconstructed using Neurolucida showing autapses in red circles. Note that all autapses are present in close distance to the soma. **F**. Pie charts depicting the percentage of cells showing (1) both structural and physiological autapses, (2) no structural nor physiological autapses and (3) only structural autapses. Note that only mice that underwent juvenile demyelination have PV interneurons which show structural autapses but no autaptic response. **G**. Representative voltage steps showing an AutIPSC response in blackand no response in red. The red trace shows a cell that had NO autaptic response. Scale:100 pA, 5ms. **H**. Shiverer mice show a clear decrease in the percentage of cells showing an autaptic response compared to control mice (*Fisher’s exact test; p = 0.0516*). **I**-**J**. There was no difference between groups in the amplitude (I) or decay (J) of the autaptic post-synaptic currents in cells that had autapses in the Shiverer mice (light red) compared to the control mice (black) (*t-test; n = 27/29 cells from 9 mice per group, p = 0.7167 (I) and p = 0.2377 (J*). **K.** Representative voltage steps showing an AutIPSC response in control (black) and adult demyelination (blue) groups. The red trace shows a cell that had NO autaptic response. Scale: 50 pA, 2 ms. **L**. Adult demyelination mice show no difference in the percentage of cells showing an autaptic response compared to control mice (*Fisher’s exact test; p = 0.7488*). **M**. There was a significant decrease in the averaged amplitude of AutIPSCs recorded in mice that underwent adult demyelination (*t-test; n = 26/18 cells from 8-6 mice per group, *p = 0.0194*). **N**. There was no difference in the decay of the autaptic post-synaptic currents (*t-test; n = 26/18 cells from 8-6 mice per group, p = 0.7804*).

Next, we went to back to the reconstructed cells and examined whether they had morphological evidence of autapses, defined as colocalization between PV interneuron axon terminals and its own soma or proximal dendrite **(Figure 7E-F**). We observed autaptic morphologies in 88.2% (15 out of 17) of control mice, but only 70.8% (17 out of 24) of PV interneurons from mice with juvenile demyelination, suggesting that an important mechanism underlying the loss of autaptic transmission following juvenile demyelination may involve overt structural loss of autapses (Fisher’s Exact Test, *\*\*p=*0.007; **Figure 7F**). In PV interneurons without an autaptic morphology, we never observed a functional autaptic response (*n*=9 cells). Furthermore, in mice with juvenile demyelination, even among PV interneurons with morphological evidence of autapses, over one-quarter showed no functional autaptic response (6 out of 24 cells in juvenile demyelination group compared to 0 out of 17 control cells, Fisher’s Exact Test for all variables, ****p<0.0001; **Figure 7E-F**). We also recorded autaptic responses from PV interneurons from *shiverer* mice. 59.5% (15 out of 37) of PV interneurons from *shiverer* mice showed no autaptic responses compared to only 16.0% (4 out of 25) in their wild-type littermates (Fisher’s Exact Test, p=0.051; **Figure 7G,H**). Amplitude and decay of the autaptic responses were also not affected (**Figure 7I,J**). Finally, to confirm that our results reflect an impairment due to the loss of myelin during development, we recorded autaptic responses from PV neurons following adult demyelination and we found no difference in the percentage of cells showing an autaptic response in mice from the adult demyelination group (65.0%; 7 out of 20) compared to control mice (72.0%; 7 out of 25) (*Fisher’s Exact Test, p=*0.749; **Figure 7K-N**).

In view of the observed impairment of sustained high-frequency firing following juvenile demyelination (**Figure 2**), we next examined whether autaptic plasticity at high frequency is also affected. Evoked autaptic responses were recorded following 100 APs at 200 Hz in voltage-clamp mode. Although control mice showed clear paired-pulse facilitation of autaptic transmission (IPSC2/1; mean paired-pulse ratio (PPR): 1.23), mice with juvenile demyelination showed no evidence of plasticity with a mean PPR of 1.02 (**Figure 8A,B)**. This impairment in autaptic plasticity was also observed in *shiverer* mice (**Figure 8C**) but not in the adult demyelination group (**Figure 8D**). Furthermore, we noted a significant increase in failure rate at 200 Hz of the subsequent evoked autaptic responses in the juvenile demyelination group where 61.1% (11 out of 18) of PV interneurons showed at least 1 failed response compared to only 13.3% (2 out of 15) in the control group (Fisher’s Exact Test, *p=0.011; **Figure 8E,F**). In *shiverer* mice 26.7 % (4 out of 15) of PV interneurons showed at least one failed response compared to only 11.7 % (2 out of 17) in the control group and in the adult demyelination group only 10.0% (1 out of 10) of the cells showed failure in autaptic responses at 200 Hz (**Figure 8G-H**).

**Figure 8.**
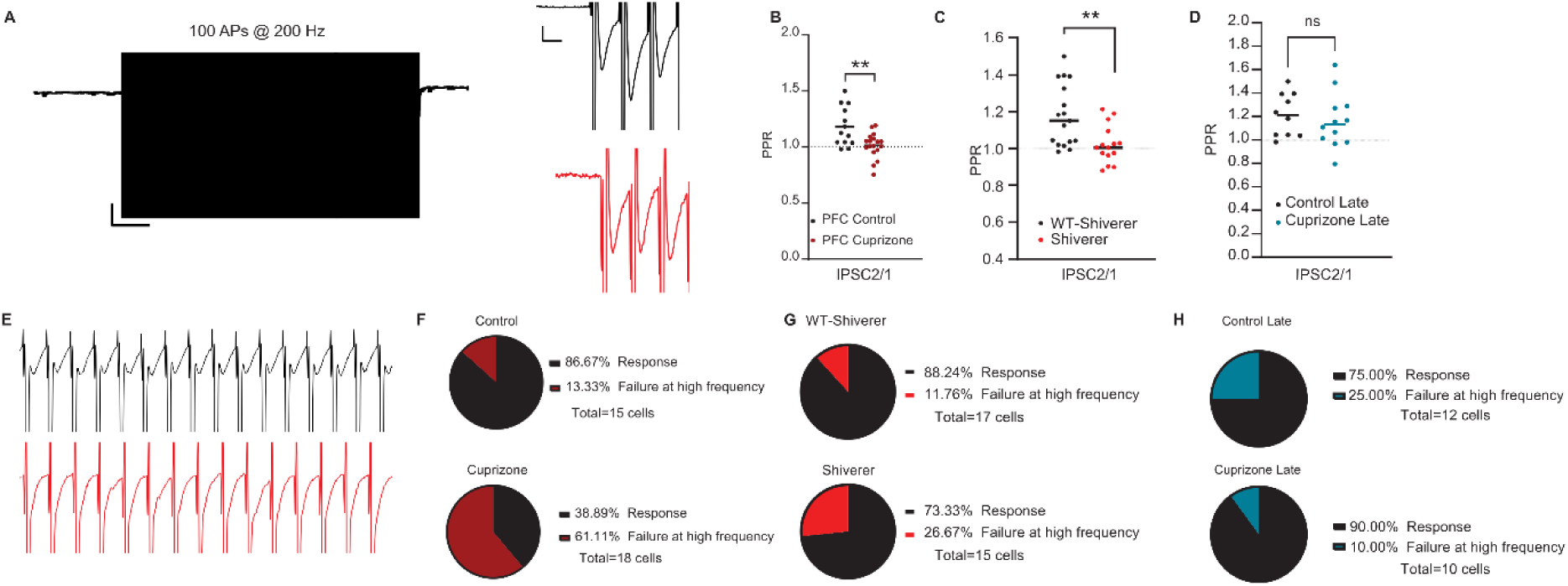
Juvenile demyelination impairs autaptic plasticity. **A.** *Left.* Example trace of the train of 100 AP at 200 Hz in voltage-clamp mode. *Right.* Inset of the first three unitary autaptic IPSCs in a train of 100 evoked at 200 Hz in PFC PV interneurons from control mice (black) and cuprizone-treated mice (red); Note the clear facilitation in the second response in the control mice. **B**. Quantification of the paired-pulse ratio (PPR) of both control (black dots) and cuprizone-treated (red dots) mice reveal an impairment in short-term plasticity at autaptic site following juvenile demyelination (*t-test: n =13,18 cells from 7-8 mice per group, **p = 0.0072*). **C**. Short-term plasticity of autaptic responses at 200 Hz is impaired in shiverer mice (light red) compared to control (black) mice (*n=17/15 cells from 9 mice per group, **p = 0.0041 (IPSC2/1)*). **D**. No difference was detected in the paired-pulse ratio (PPR) between control (black dots) and adult demyelination (blue dots) (*n= 12,10 cells from 8-6 mice per group, p = 0.5780 (IPSC2/1)*). **E.** Example traces of the first 15 unitary autaptic IPSCs in a train of 100 evoked at 200 Hz in PFC PV interneurons from control mice (black) and cuprizone-treated mice (red) ; Note that many evoked responses failed in the cuprizone mice . **F**. Cuprizone-treated mice show a significant increase in the percentage of cells showing a failed autaptic response at 200 Hz (*Fisher’s exact test; *p = 0.0110*). **G**. There was no significant difference in the percentage of cells showing a failed autaptic response at 200 Hz remains in the Shiverer group compared to the control group (*Fisher’s exact test; p = 0.3828*). **H**. There was no difference in the percentage of cells showing a failed autaptic response at 200 Hz remains in the adult demyelination group compared to the control group (*Fisher’s exact test; *p = 0.5940*).

#### Loss of PV interneuron autapses predicts high-frequency firing impairments

Juvenile demyelination causes impairment in high-frequency firing of PV interneurons and a loss of functional autapses, however, is there a link between these two phenotypes? To explore this further, we divided the cells from both the juvenile demyelination group and the control group into (1) cells with an autaptic response and (2) cells with *no* autaptic response (**Supplementary Figure 5**). In the control group, we found that PV interneurons with an autaptic response were less excitable than cells with no autaptic response, with a difference in rheobase of ∼40 pA (**Supplementary Figure 5A,B**). PV interneurons with autapses also showed an increased AHP without a change in AP width (**Supplementary Figure 5C,D**). Importantly, although cells with no autapses had an increased firing rate following low-current injections (0-250 pA), their firing rate was attenuated in response to high-current injections (300-600 pA) compared to cells with autapses (**Supplementary Figure 5E,F**). These data indicate that autapses are necessary but not sufficient for high-frequency firing in PV interneurons. How was this different in the juvenile demyelination group? In mice that underwent juvenile demyelination, 60.0% (18 out of 30) of PV interneurons that had no autaptic response showed impaired firing at high frequencies (100-250 Hz). Conversely, 60.0% (12 out of 20) of PV interneurons with impaired firing at high frequencies had no autaptic response (**Supplementary Figure 5G-H**), suggesting that the absence of autapses in the juvenile demyelination group increased the likelihood of PV interneurons exhibiting impaired firing. Indeed, 31.0% (16 out of 52) of all PV interneurons from the juvenile demyelination group showed both no autaptic responses and an impaired firing phenotype versus only 4.6% (2 out of 43) in the control group. While these results indicate that the relationship between the loss of functional autapses and the impairment in high-frequency firing in the juvenile demyelination group may not be directly causal, they indicate that a loss of autapses during the critical period of development of PV interneurons might facilitate their probability of failure to fire at high frequencies.

### Can we rescue PV neuron firing properties with remyelination in adulthood?

#### Remyelination in adulthood only partially restores PV interneuron properties in the PFC

Our data reveal a clear impairment in PV interneuron properties following juvenile demyelination. We next questioned whether remyelination could restore PV interneuron function in PFC. To that end, mice underwent juvenile demyelination from P21 to P60, after which they returned to a normal diet from P60 to P100 (**Figure 9A-B**). We then performed whole-cell recordings from PV interneurons in the PFC. Both the input resistance and the sag amplitude were restored to control levels upon remyelination (**Figure 9C-H**). Interestingly, we found that remyelination caused a significant increase in the rheobase of PV interneurons, promoting a decreased excitability of these cells. When examining the AP waveform, we detected a small increase in the decay of the AP in the remyelination group, even though AP half-width was not significantly different (**Figure 9I-M**). PV interneurons from the remyelination group still showed decreased firing frequency in response to increased current injections (**Figure 9N-P**). Even after remyelination, 26.2% (11 out of 42) of the cells still could not sustain their firing at high frequency, compared to 5.0 % (1 out of 20) in control mice (Fisher’s Exact Test, p=0.083; **Figure 9Q,R**). We then investigated whether self-inhibitory transmission was restored by remyelination (**Figure 9S-X**). Remyelination resulted in a rescue of autaptic neurotransmission (autaptic responses: remyelination, 68.4%, 19 out of 28; control, 75.0%, 15 out of 20; *Fisher’s Exact Test, p=*0.764;) (**Figure 9S-T**). The amplitude and decay of the responses were also similar across groups (**Figure 9U,V**). No significant difference between groups was detected in short-term plasticity at 200 Hz (IPSC2/1; mean PPR: 1.18 for remyelination group and 1.12 for control group) (**Figure 9W**). Yet, there was a significant difference in the percentage of cells that showed failures at 200 Hz in the remyelination group (Fisher’s Exact Test, *p=0.023; **Figure 9X**). These results suggest that remyelination leads to a partial rescue of PFC PV interneurons, in which the impairment remains apparent when PV interneurons are driven to fire at high frequencies.

**Figure 9.**
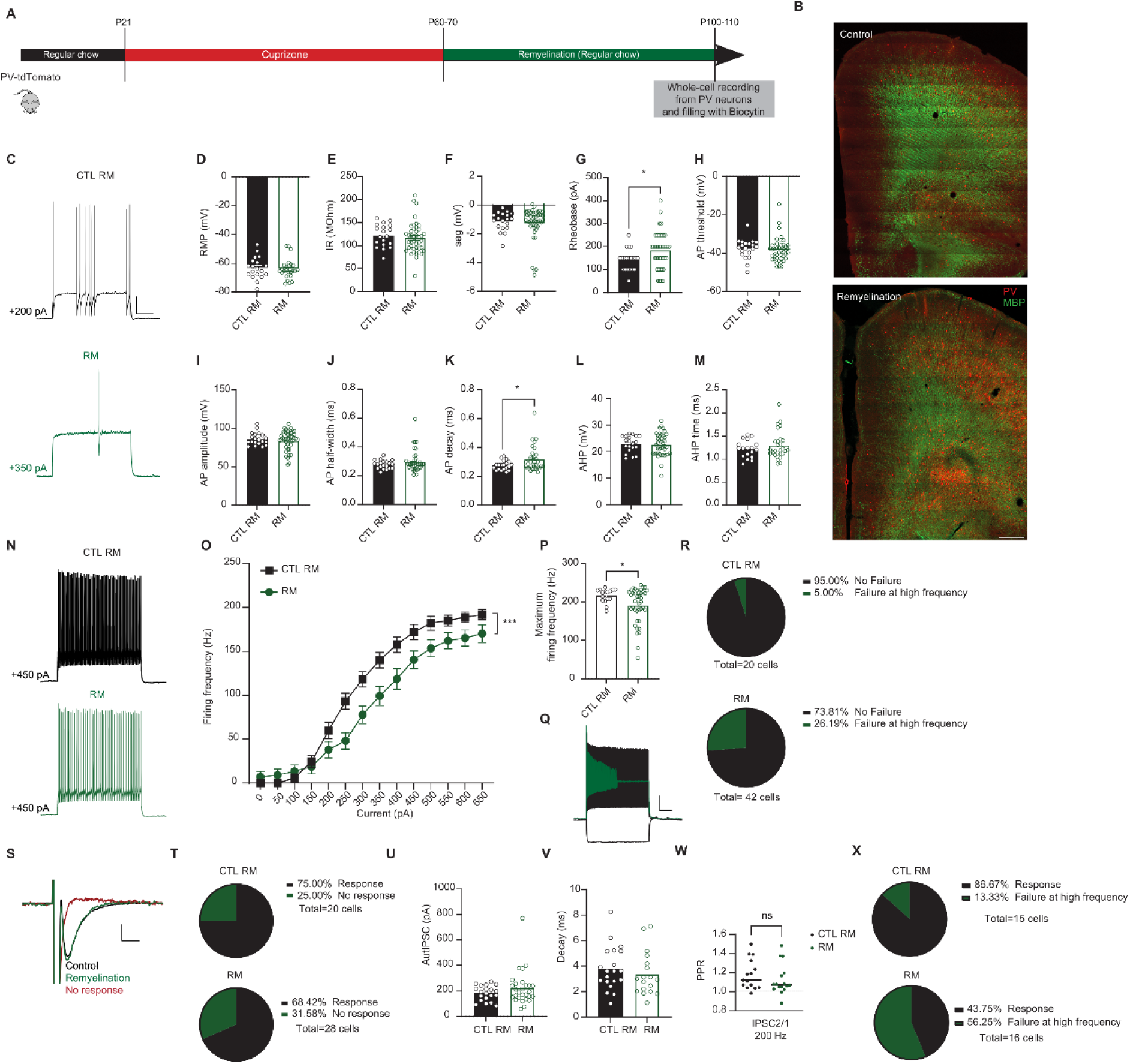
Remyelination in adulthood leads to an incomplete restoration of PV interneuron properties. **A.** Experimental design for remyelination in PV-tdTomato mice **B**. *Upper panel.* Confocal overview image of PFC in PV-tdTomato animal (tdTomato+, red) overlaid with myelin basic protein (MBP, green). *Lower panel.* Confocal overview image immunolabeled for MBP expression showing remyelination of the PFC region following 5 weeks of normal diet. Sacle: 300 um. **C**. Representative traces of voltage responses following a depolarizing step from control (black) and remyelination (green) mice illustrating an increased in Rheobase in mice after remyelination. Scale: 20 mV, 100ms. **D**-**H**. Summary data showing the averaged (± s.e.m) of the following intrinsic properties: (D) Resting membrane potential (RMP) (*t-test; n = 25/19 cells from 8 mice per group, p = 0.7435*), (E) Input resistance (IR) (*t-test; n = 25/19 cells from 8 mice per group, p = 0.5626*), (F) Sag (*t-test; n = 25/19 cells from 8 mice per group, p = 0.6088*), (G) Rheobase (*t-test; n = 25/19 cells from 8 mice per group, *p < 0.05*) and (H) Action potential (AP) threshold (*t-test; n = 25/19 cells from 8 mice per group, p = 0.6472*). **I**-**M**. Summary data showing the averaged (± s.e.m) of the following AP waveform properties: (I) AP amplitude (*t-test; n = 25/19 cells from 8 mice per group, p = 0.4951*), (J) AP half-width (*t-test; n = 25/19 cells from 8 mice per group, p = 0.2202*), (K) AP decay (*t-test; n = 25/19 cells from 8 mice per group, *p < 0.05*), (L) After-hyperpolarization (AHP) amplitude (*t-test; n = 25/19 cells from 8 mice per group, p = 0.8156*) and (M) AHP time (*t-test; n = 25/19 cells from 8 mice per group, p = 0.3123*). **N**. Representative traces of voltage responses following +450 pA current injection. **O**. Average action potential (AP) frequency in response to 0-650 pA current steps illustrating a significant decrease in PV interneuron firing frequency in remyelination (green) mice (*group x current two-way repeated measures: n = 25/19 cells from 8 mice per group: F(13,546)= 3.962, ***p < 0.001)*. **P**. Summary data of the maximum firing frequency per group (*t-test; n = 25/19 cells from 8 mice per group, *p < 0.05*). **Q**. Example trace of a PV interneuron that is unable to maintain high-frequency firing at increased current injection. **R**. Percentage of cells that failed to maintain high frequency firing at > 500 pA is still not fully recovered after remyelination (*Fisher’s exact test; p = 0.0828*). **S**. Representative voltage steps showing an AutIPSC response in control (black) and remyelination (green) groups. The red trace shows a cell that had NO autaptic response. Scale: :50 pA, 2ms. **T**. Remyelination mice show no difference in the percentage of cells showing an autaptic response compared to control mice (*Fisher’s exact test; p = 0.7639*). **U-V**. There was no difference between groups in the amplitude (U) or decay (V) of the autaptic post-synaptic currents in cells that had autapses in the remyelination group (green) compared to the control group (black) (*t-test; n = 28/20 cells from 8 mice per group, p = 0.1939 (U) and p = 0.4328 (V)*). **W-X**. Paired-pulse ratio (PPR) of both control (black dots) and remyelination (green dots) mice shows no difference in short-term plasticity at autaptic sites specifically at 200 Hz (*t-test; n = 15/16 cells from 8 mice per group, p = 0.3388*). Yet, the percentage of cells showing a failed autaptic response at 200 Hz remains increased in the remyelination group compared to the control group (*Fisher’s exact test; *p = 0.0233*).

#### Juvenile demyelination has long-lasting effects on social behavior

As previous studies have demonstrated that impaired myelination during this critical period leads to impaired social behavior in adulthood (Makinodan et al. 2012), we next explored whether mice that underwent a period of juvenile demyelination followed by remyelination, have any deficits in social behavior (**Figure 10**). More specifically, we were interested in examining whether the impairments in PV interneuron properties at high frequencies were accompanied with behavioral changes. To that end, we performed a social behavior test, the three-chamber test, which is a commonly used behavioral assay in mice to assess social behavior in mice modelling neurodevelopmental disorders like schizophrenia. Furthermore, mice with PFC lesions or dysfunction show reduced interaction with a novel mouse, indicating a lack of sociability. In the first phase of the test, sociability was assessed (preference of mouse over object). Mice that had undergone juvenile demyelination followed by a period of remyelination showed no preference towards the mouse compared to the object, while control mice spent significantly more time exploring the mouse compared to the object (**Figure 10B**). In the second phase, social novelty preference was tested (preference of novel mouse over familiar mouse). Both groups spent significantly more time exploring the novel mouse compared to the familiar mouse (**Figure 10C**). These data indicate that transient juvenile demyelination results in long-lasting selective impairments of sociability during adulthood.

**Figure 10.**
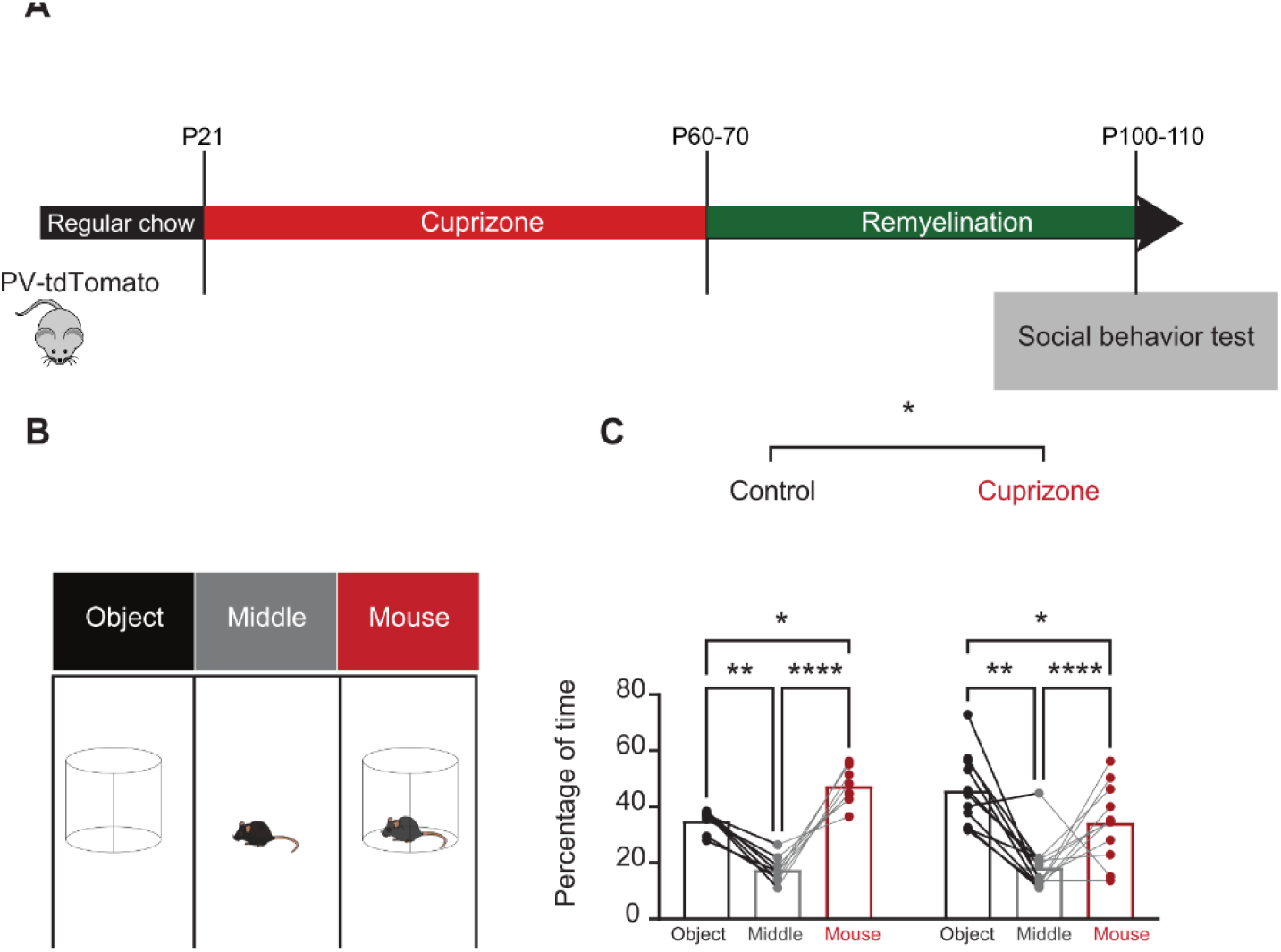
Mice that underwent juvenile demyelination present a significant deficit in social behavior in adulthood. **A.** Experimental design. **B**. Behavioral results for sociability show that cuprizone mice do not show a social preference while control mice interacted significantly more with the mouse compared to the object (*n= 8 control mice and 11 cuprizone mice; two-way repeated measures ANOVA, group x stimulus interaction F(2,34) =4.531, *p = 0.0180 followed by post hoc LSD test, *p < 0.05, **p < 0.01, ***p < 0.001*). **C.** Experimental design for the second phase of the three-chamber test. **B**. Behavioral results for social novelty show that both cuprizone mice and control mice showed a preference to interact with the novel mouse compared to the familiar mouse (*n= 8 control mice and 11 cuprizone mice; two-way repeated measures ANOVA, group x stimulus interaction F(2,32) =0.2412, p = 0.7871 followed by post hoc LSD test, *p < 0.05, **p < 0.01, ***p < 0.001*).

## Discussion

### Myelination of PFC PV interneurons during adolescence is crucial for their structural and electrophysiological maturation

Several studies have highlighted the importance of adult PV interneuron myelination in proper PV interneuron function, such as inhibitory control of local oscillations, experience-dependent plasticity or AP conduction velocity increase along PV axons (Micheva et al. 2021a; Dubey et al. 2022; S. M. Yang et al. 2020). Yet, little is known about whether PV interneuron myelination is important for cell autonomous development and maturation. Given the implications of PFC development in many neuropsychiatric disorders, along with a clear role of PV interneuron dysfunction in such disorders, adding to it the various reports of myelin deficits in patients with neuropsychiatric disorders (J. Stedehouder and Kushner 2017), we aimed to investigate whether an early disruption of PV interneuron myelination, during the critical period of PFC development, can lead to an impairment in cell autonomous maturation and development. Many studies report an extended critical period for PFC maturation from P21 to P35 (Toritsuka, Makinodan, and Kishimoto 2015; Makinodan et al. 2012; 2009; Bicks et al. 2020). We used the cuprizone model starting at P21 to induce juvenile demyelination. Our data revealed that juvenile demyelination leads to persistent alterations in PV interneuron morphology in adulthood. Specifically, we found a decrease in total axonal length, as well as the number of axonal branches among PV interneurons from mice that underwent juvenile demyelination. Axons of PV interneurons have previously been reported to have considerable morphological plasticity during the second and third postnatal week, at a time where myelination is starting to peak (Micheva et al. 2021b; Y. Yang et al. 2016; Jeffrey Stedehouder et al. 2018). To what extent might these structural changes be related to PV interneuron intrinsic properties and electrophysiological development? Our data revealed that PV interneurons in the PFC of mice that underwent juvenile demyelination show clear maturation deficits in their intrinsic properties. Specifically, we observed an increase in the input resistance and sag amplitude, an increase in AP width and a substantial decrease in the firing frequency, especially at high frequencies, all reminiscent of immature PV interneurons (Ethan M. Goldberg et al. 2011; Miyamae et al. 2017). Notably, among mice that underwent juvenile demyelination, ∼40% of the PV interneurons were unable to sustain high-frequency firing, for which their maximum firing rate was substantially decreased. Similar observations were made in the *shiverer* model, suggesting that the lack of myelin during this critical period of development had significant impact on the electrophysiological maturation of these cells. Indeed, in mice that underwent adult demyelination, while PV interneurons were less excitable, which is in agreement with previous studies (Dubey et al. 2022), high-frequency firing was not affected. Taken together, our results indicate that the ontogeny of myelination is crucial for the proper structural and physiological development of PV interneuron.

### Kv3 is crucial for PV maturation and myelination

PV interneuron electrophysiological maturation has been closely linked to the development of distinct ion channels that allow for their unique ability to fire at high frequencies. Specifically, Kv3 channels have been shown to double their expression level from P18 to P30 (Ethan M. Goldberg et al. 2011). Therefore, we next explored whether a loss of myelin during adolescence could be affecting Kv3 expression and activity levels in PV interneurons of the PFC. Previous studies have demonstrated that myelination causes alterations in Na+ and K+ channels in pyramidal neurons (Hamada and Kole 2015). Our results confirmed a clear decrease in Kv3.1 expression by immunohistochemistry, and K^+^ current using voltage-clamp recordings from PV interneurons of the PFC in mice that underwent juvenile demyelination. We next tested whether AUT00201, a positive modulator of Kv3 K+ currents, could restore Kv3 currents in PV interneurons from mice that underwent juvenile demyelination. Bath application of AUT00201 rescued the deficit in AP width but not the alteration of high-frequency firing. This indicates that, while acute Kv3 activation can restore AP waveforms in these cells, the loss of Kv3 expression due to juvenile demyelination likely requires a more chronic intervention to fully rescue PV maturation. Moreover, it is not unlikely that other channels might also be affected by juvenile demyelination. In particular, the changes in the sag we observed following juvenile demyelination and in *shiverer* mice are reminiscent of alterations in HCN channel function. HCN channels have been reported to be exclusively localized on axons in PV interneurons, and play a critical role in enhancing AP initiation during sustained high-frequency firing and in increasing the velocity by which APs propagate along PV interneuron axons (Roth and Hu 2020). Moreover, expression of TASK-1 and -3 channels, which are critical for maintaining the RMP and modulation of the excitability of PV interneurons, increase during adolescence, for which PV interneurons from TASK-1/3 double knock-out mice exhibit increased input resistance, wider AP and a decreased firing rate (E. M. Goldberg 2005).

### Myelination of PFC PV interneurons during adolescence is important for self-inhibitory transmission

What happens to autapses following juvenile demyelination and the consequent aberrant development of PV interneurons? Our data revealed a significant decrease in the number of PV interneurons that have autaptic responses after juvenile demyelination, but not after adult demyelination. In examining reconstructed neurons, we found that ∼25% of PV interneurons from the juvenile demyelination group exhibited structural autapses but no autaptic response, indicating that the loss of myelin during adolescence led to an impairment of autaptic neurotransmission. Moreover, ∼30% of the cells had no discernible autapses, compared to only 11% in controls, suggesting a complete loss of autapses in some PV interneurons due to juvenile demyelination. Yet, more than 45% of PV interneurons of mice that underwent juvenile demyelination still had functional autapses, in which the amplitude of the responses appeared largely unaffected. Therefore, we next investigated whether plasticity at autaptic sites was impaired as a result of juvenile demyelination. Our data uncovered an impairment in short-term plasticity at autaptic sites at 200 Hz, indicating that myelination during adolescence is required for adult synaptic plasticity at high frequencies. Results in *shiverer* PV interneurons confirmed these data. Conversely, in mice that underwent adult demyelination, short-term plasticity at autaptic sites was not affected, although, the amplitude of the inhibitory autaptic response was decreased. This is likely due to a deficit in inhibitory synapse maintenance caused by the loss of myelin, as previously reported (Dubey et al. 2022). Together, our data suggest that juvenile demyelination can cause a loss of functional autapses in PV interneurons of the PFC and impair autaptic neurotransmission. It remains to be investigated whether the deficits in short-term plasticity recorded in PV interneurons from mice that underwent juvenile demyelination are due to failures in AP-triggered GABA release or failures of AP propagation along demyelinated axons.

### Autapses play a role in high frequency firing of PV interneurons

Previous studies have demonstrated that fast-spiking interneurons exhibit autapses, which are synaptic contacts between the cell’s own soma or dendrite and its own axon (Micheva et al. 2021b; Deleuze et al. 2019; Deleuze, Pazienti, and Bacci 2014; Bacci, Huguenard, and Prince 2003; Szegedi et al. 2020). While there is still a lot unknown about the role of these autapses, it has been proposed that they play an important role in the temporal control of microcircuits by modulating PV interneuron excitability. Specifically, autaptic neurotransmission has been proposed as a particular form of cortical disinhibition regulating PV interneuron influences on network oscillations (Deleuze et al. 2019; Deleuze, Pazienti, and Bacci 2014). Approximately 70 to 80% of PV interneurons have autapses, while non-fast-spiking interneurons do not exhibit autapses (Deleuze, Pazienti, and Bacci 2014; Bacci, Huguenard, and Prince 2003; Szegedi et al. 2020; Connelly and Lees 2010). PV interneuron autapses are GABAergic with high probabilities of release, in which plasticity appears to follow largely similar rules as non-autaptic inhibitory postsynaptic sites (Deleuze et al. 2019; Bacci, Huguenard, and Prince 2003; Szegedi et al. 2020; Connelly and Lees 2010). Interestingly, a recent study found that autaptic neurotransmission induced larger responses than other synaptic inputs in a subset of PV interneurons and they estimated that autaptic transmission accounted for approximately 40% of the global inhibition that PV interneurons received (Deleuze et al. 2019).

In the current study, we aimed to explore whether autapses might play a role in PV interneuron’s ability to fire at high frequency, which in turn has been demonstrated to be of importance for network oscillations and higher cognitive processing (Deleuze et al. 2019; Deleuze, Pazienti, and Bacci 2014). We show that PV interneurons without autapses are more excitable, displaying a lower rheobase and a change in the AP’s AHP. Interestingly, these cells had higher firing frequencies at low current injections, yet they had a significantly decreased maximum firing rate. These data suggest that autapses contribute to PV interneuron’s ability to fire at high frequency by controlling the excitability and the timing of PV interneurons spiking activity, as proposed in previous studies (Deleuze, Pazienti, and Bacci 2014; Bacci and Huguenard 2006; Szegedi et al. 2020).

### Remyelination after juvenile demyelination is unable to fully rescue PV interneuron function and PFC-dependent social behavior

Remyelination studies have revealed that in only 4 weeks after prolonged cuprizone treatment, nearly complete remyelination can occur (Zendedel, Beyer, and Kipp 2013; Gingele et al. 2020; Hiremath et al. 1998; ‘De- and Remyelination in the CNS White and Grey Matter Induced by Cuprizone: The Old, the New, and the Unexpected’ 2011). We therefore investigated whether remyelination could restore PV interneuron properties and autaptic neurotransmission following juvenile demyelination. Our data showed that while many of the membrane properties were rescued by remyelination, PV interneuron firing rate was still decreased after remyelination and more than 25% of the cells could not sustain their firing rate at high frequencies. Interestingly, autaptic responses at high frequency were also still impaired. This suggests that restoring myelin in the PFC might have reinstated some channel properties in PV interneurons, yet the ability of PV interneurons to fire at high frequency was not fully restored. We propose that the loss of myelin during this critical period of PFC development constrict any full rescue of PV interneuron properties. This is in line with many previous works that have confirmed that this is a sensitive period for PFC development and any intervention during this vulnerable period could have long-lasting effects (Hensch 2005; Makinodan et al. 2012; 2009; Chini and Hanganu-Opatz 2021).

Studies have shown that insults to the PFC during the juvenile critical period can have long-lasting detrimental effects on social behavior (Klune, Jin, and DeNardo 2021; Toritsuka, Makinodan, and Kishimoto 2015). Moreover, impairments in PV interneuron during this juvenile period lead to deficits in adult social behavior (Fang et al. 2022; Bicks et al. 2020). Accordingly, we uncovered that mice which underwent juvenile demyelination followed by a period of remyelination, a timepoint where we showed that PV interneurons function was not yet fully restored in the PFC, exhibited social deficits. This data implies that juvenile demyelination can lead to long-last impairment in PV interneuron function which could explain social impairments in adulthood.

### Relevance for human disorders

The development of PFC circuits is known to extend beyond the juvenile developmental period (Klune, Jin, and DeNardo 2021; Bitzenhofer et al. 2021). It is characterized by an essential maturation of inhibitory networks and the establishment of excitatory/inhibitory balance, which in turn give rise to various brain oscillations involved in higher cognitive functions (Bitzenhofer et al. 2021; Bitzenhofer, Pöpplau, and Hanganu-Opatz 2020). PV interneuron development follows a prolonged period of maturation that also extends into adolescence (Ethan M. Goldberg et al. 2011; J. Stedehouder and Kushner 2017; Miyamae et al. 2017). While many studies have attributed an impairment in PFC development and PV interneuron maturation to many neuropsychiatric disorders (van Os and Kapur 2009), little is known about whether there is a link between myelination and PV interneuron development in the PFC. Makinodan and colleagues were the first to identify a critical period for oligodendrocyte maturation and myelination for PFC-dependent social behaviour (Makinodan et al. 2012). They showed that social isolation of mice between P21 and P35 can have long lasting effects on PFC myelination, where mice that underwent social isolation in this period showed less myelin expression in the PFC compared to mice that were socially isolated after P35 and control mice. Furthermore, mice in which myelination was impaired because of loss of ErbB3 signalling showed similar social behaviour deficits to the ones elicited by social isolation during the critical period, confirming that myelination of the PFC during this period is necessary for normal social behaviour in the adult. Interestingly, another study uncovered that demyelination in juvenile mice, but not in adulthood, can lead to long-lasting impairments in social interaction in mice (Makinodan et al. 2009). Finally, it has been shown that social isolation during this critical period (P21-35) can have significant effects on PV interneuron activity in the PFC, specifically PV interneurons had a decreased firing rate. Taken together, we propose that our data highlights an important link between PV interneuron myelination and their development in the PFC during this critical period, which offers key insights into the mechanisms underlying the pathophysiology and potential therapies for neurodevelopmental disorders.

## Materials and Methods

All experiments were conducted under the approval of the Dutch Ethical Committee and in accordance with the Institutional Animal Care and Use Committee (IACUC) guidelines. The Pv-tdTomato [Tg(Pvalb-tdTomato)15Gfng] line was used as well as the Shiverer [B6;C3Fe.SWV-Mbpshi /J (Shiv) (Chernoff 1981; www.jax.org/strain/001428)] line that was crossed with the Pv-tdTomato line .

For cuprizone treatment, mice received 0.2% (w/w) cuprizone (Bis(cyclohexanone)oxaldihydrazone (C9012, Merck) added to grinded powder food or to food pellets (Envigo). For the juvenile demyelination group, mice received fresh cuprizone food starting at P21 for a period of 6-7 weeks while control mice received control food, both *ad libitum*. For the adult demyelination group, mice received either cuprizone food (cuprizone group) or normal chow food (control group) starting P60 for a period of 6-7 weeks. The average maximum weight loss during cuprizone treatment was around 15 % in the adult cuprizone group and around 7 % in the juvenile demyelination group. All mice from both sexes were used for these experiments. All mice were maintained on a regular 12 h light/dark cycle at 22 °C (±2 °C) with access to food and water *ad libitum*.

### Slice preparation and electrophysiology

Whole-cell recordings of PV interneurons in layers 2-3 in the PFC were performed as described previously (J. Stedehouder et al. 2017; Pascual-García et al. 2024). Briefly, after decapitation, brains were placed in ice-cold partial sucrose-based solution containing (in mM): sucrose 70, NaCl 70, NaHCO3 25, KCl 2.5, NaH2PO4 1.25, CaCl2 1, MgSO4 5, sodium ascorbate 1, sodium pyruvate 3, and D(+)-glucose 25 (carboxygenated with 5% CO2/95% O2). Coronal slices from the prefrontal cortex (300 μm thick) were obtained with a vibrating slicer (Microm HM 650V, Thermo Scientific) and incubated for 45 min at 34°C in holding artificial cerebrospinal fluid (ACSF) containing (in mM): 127 NaCl, 25 NaHCO3, 25 D(+)-glucose, 2.5 KCl, 1.25 NaH2PO4, 1.5 MgSO4, 1.6 CaCl2, 3 sodium pyruvate, 1 sodium ascorbate, and 1 MgCl2 (carboxygenated with 5% CO2/95% O2). Next, the slices recovered at room temperature for another 15 min. Slices were then transferred into the recording chamber where they were continuously perfused with recording ACSF (in mM): 127 NaCl, 25 NaHCO3, 25 D-glucose, 2.5 KCl, 1.25 NaH2PO4, 1.5 MgSO4, and 1.6 CaCl2. Cells were visualized using an upright microscope (BX51WI, Olympus Nederland) equipped with oblique illumination optics (WI-OBCD; numerical aperture 0.8) and a 40× water-immersion objective. Images were collected by a CCD camera (CoolSMAP EZ, Photometrics) regulated by Prairie View Imaging software (Bruker). Layer II-III pyramidal cells in the somatosensory cortex were identifiable by their location and morphology. Electrophysiological recordings were acquired using HEKA EPC10 quattro amplifiers and Patchmaster software (10 Hz sampling rate) at 33°C. Patch pipettes were pulled from borosilicate glass (Warner instruments) with an open tip of 3.5–5 MegaOhm of resistance and filled with intracellular solution containing (in mM) 125 K-gluconate, 10 NaCl, 2 Mg-ATP, 0.2 EGTA, 0.3 Na-GTP, 10 HEPES and 10 K2-phosphocreatine, pH 7.4, adjusted with KOH (280 mOsmol/kg), with 5 mg/mL biocytin to fill the cells. Series resistance was kept under 20 M with correct bridge balance and capacitance fully compensated; cells that exceeded this value were not included in the study. Cells were filled with biocytin for at least 20 min. .AMPA-mediated currents were blocked using DNQX (10 μM,, HelloBio). Gabazine (10 μM, HelloBio) was used to block GABAARs. AutIPSC recordings were measured as described previously. For Kv current recordings, holding potential was -70 mV and voltage steps from -70 mV to +60 mV (in 10 mV increments) were recorded to extract high voltage activated potassium currents. Tetrodotoxin (1uM) was used to suppress sodium-activated channels. TEA (1mM) was applied to the bath following baseline recordings. The resulting voltage currents were obtained by subtraction of baseline from TEA-sensitive currents. Leak currents were removed online.

Intrinsic passive and active membrane properties were recorded in current-clamp mode, both at resting membrane potential and at -70 mV, by injecting 500-ms of increasing current stimuli from - 300 pA to +650 pA, at intervals of 50 pA. Data analysis was conducted using a custom-designed script in Igor Pro-9.0 (Wavemetrics).

### AUT00201 compound

AUT00201 (Autifony Therapeutics) is a small molecule modulator of Kv3.1 and Kv3.2 channels. AUT00201 (10 mM stock solution) was dissolved in DMSO (0.1%), and the final concentration used in the recording chamber was 1 uM.

### Immunohistochemistry

Patch-clamp recorded PV interneurons were filled with 5mg/mL biocytin during whole-cell recordings and then fixed with 4% (PFA) overnight and stored in PBS at 4°C. Stainings were done as previously described (Stedehouder et al. 2019). The following primary antibodies were used: rabbit anti-Cherry (1:500, Millipore, Abcam, ab167453), mouse anti-PV (1:1000, Swant, 235), and goat anti-MBP (1:300, Santa Cruz, C-16, sc-13 914).

### Confocal imaging and reconstruction

Images were taken using a Zeiss LSM 700 microscope. Biocytin-filled whole-cell overview image was acquired using 63x magnification objective with tiled *z*-stack images (512 × 512 pixels) with a step size of 1 μm. Three-dimensional reconstruction of the cells was performed using Neurolucida software (MBF Bioscience) as previously described (Stedehouder et al. 2017; Stedehouder et al. 2019). Autapses were defined as individual sites where the distance between an axon and a dendrite of the same cell was less than 1 μm.

### Social Behavior

The three chambers test was performed in the sociability cage. Mice were first habituated for 5 min in the middle chamber and then 10 min with access to all 3 chambers empty, one day prior to testing. For the sociability test, the test mouse was first placed in the center chamber for 5 min with the doors closed. Then the doors were opened while a mouse of similar age and same sex was kept under a wire cage in one of the side chambers (novel mouse). The other chamber contained an empty wire cage (novel object). The test mouse was left to freely explore the 3 chambers for 10 minutes. For the social novelty test, the test mouse was then placed back in the center chamber and the doors were closed again. A novel mouse was placed in the empty wire cage. The doors were then opened and the test mouse was again allowed to freely explore both mice for 10 min. Behavior was recorded and scored by Ethovision (Noldus: v.9,14). The time spent in each chamber was extracted and the percentage of time spent in the chamber was calculated as the total time spent in that chamber divided by the total time spent in all 3 chambers.

### Statistical analysis

All statistical analysis were operated using GraphPad Prism 8. First, data were tested for normality. Data sets following normal distribution were analyzed using unpaired two-tailed *t*-test. Data sets without a normal distribution were analyzed using Mann-Whitney test. One-way ANOVA with LSD test for post-hoc analysis was used for group comparison for groups with equal variances. Two-way repeated measures ANOVA was used for assessing effects within groups and between groups in experiments with repeated measurements in the same cell. All quantitative data are represented as means ± standard errors of the means (s.e.m). To reduce selection bias, all mice were randomly allocated to the different groups.

## Acknowledgments

This work was funded in part by European Research Area Network ERA-NET NEURON JTC2018-024 / NWO 013.18.002 (M.P.G., S.A.K.) and ERA-PerMed2018-127 / ZonMw 456008003 (S.A.K.). We thank Nadia Pilati, Martin Gunthorpe, Martin Pue and Charles H. Large from Autifony S.r.l. for providing us with the Kv3 modulator AUT00201 and for their critical input on the subject; D. Slump for her assistance with breeding and genotyping of mice; D. Rotaru for her helpful suggestions; and all members of the Kushner laboratory for their support.

## Author contributions

S.H. and S.K. contributed to the study design. S.H, M.P.G and Y.N contributed to data collection. S.H. carried out the data analysis. S.H., M.P.G., and S.K. wrote the paper. All authors discussed and commented on the manuscript.

## Supplementary Figures

**Supp. Figure 1.**
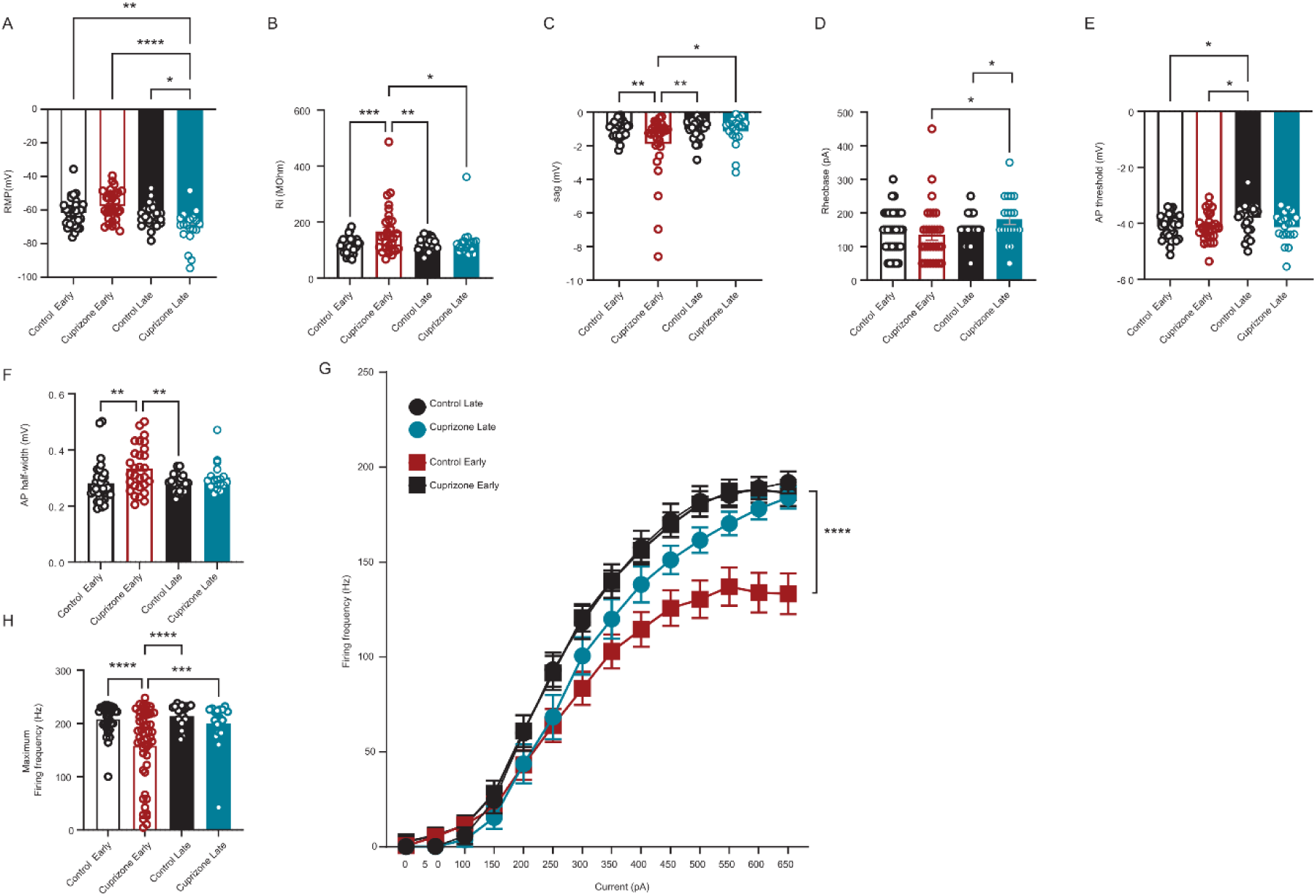
**Comparison between juvenile and adult demyelination. A-I**. Juvenile demyelination leads to impairment in PV interneuron maturation whereas adult demyelination induces a decrease in the excitability of PV interneurons. Summary data showing the averaged (± s.e.m) of the following intrinsic properties: (A) Resting membrane potential (RMP) (*ANOVA: n =36,30,19,24 cells from 9/8/8/6 mice per group, ****p < 0.0001, post-hoc LSD test: *p < 0.05, **p < 0.01, ****p < 0.0001*), (B) Input resistance (IR) (*ANOVA: n =36,30,19,24 cells from 9/8/8/6 mice per group, **p = 0.0024, post-hoc LSD test: *p < 0.05, **p < 0.01, ***p < 0.001*), (C) Sag (*ANOVA: n =36,30,19,24 cells from 9/8/8/6 mice per group, **p = 0.0083, post-hoc LSD test: *p < 0.05, **p < 0.01*), (D) Rheobase (*ANOVA: n =36,30,19,24 cells from 9/8/8/6 mice per group, p = 0.1230, post-hoc LSD test: *p < 0.05*) and (E) Action potential (AP) threshold (*ANOVA: n =36,30,19,24 cells from 9/8/8/6 mice per group, *p = 0.0452, post-hoc LSD test: *p < 0.05*), (F) AP half-width (*ANOVA: n =36,30,19,24 cells from 9/8/8/6 mice per group, **p = 0.0058, post-hoc LSD test: **p < 0.01*), (G) Average action potential (AP) frequency in response to 0-650 pA current steps illustrating no significant change in S1 PV interneuron firing frequency following juvenile demyelination (*group x current two-way repeated measures: n = 36,30,19,24 cells from 9/8/8/6 mice per group: F(39,1291)= 2.487, ****p < 0.0001*) and (H) Maximum firing frequency per group (*ANOVA: n =36,30,19,24 cells from 9/8/8/6 mice per group, ****p < 0.0001, post-hoc LSD test: ***p < 0.001, ****p < 0.0001*).

**Supp. Figure 2.**
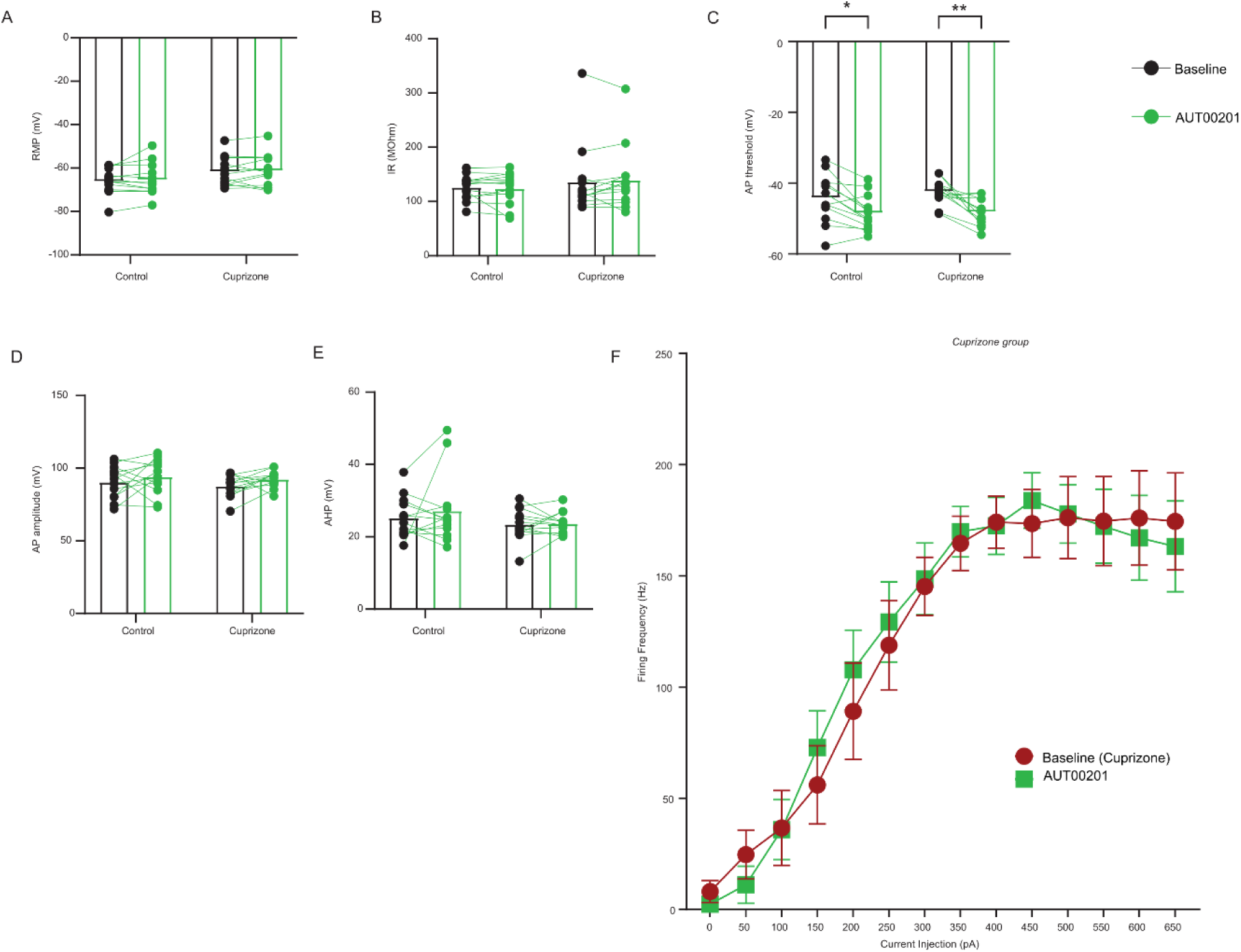
Effect of AUT00201 (1uM) on PV properties after juvenile demyelination. A-B. Bath application of AUT00201 (1uM) had no significant effect on the resting membrane potential (A) or the input resistance (B) of PV interneurons from cuprizone mice and control mice (RMP: *group x AUT00201 two-way repeated measures: n = 12,13 cells from 4 mice per group: p = 0.8593; AUT00201 effect: p = 0.2171.* IR*: group x AUT00201: p = 0.8605; AUT00201 effect: p = 0.4704*). **C**-**E**. AUT00201 decreased the AP threshold of PV interneurons in both groups (*group x AUT00201 two-way repeated measures: n = 12,13 cells from 4 mice per group: p = 0.4503; AUT00201 effect: ****p < 0.0001. Post-hoc LSD analysis: *p < 0.05, **p < 0.01*) while not having an effect on either the AP amplitude (D) or the AHP (E) (D: *group x AUT00201 two-way repeated measures: n = 12,13 cells from 4 mice per group: p = 0.8235; AUT00201 effect: p = 0.0763.* E*: group x AUT00201: p = 0.6088; AUT00201 effect: p = 0.3628*. **F**. There was no effect detected on the overall firing frequency of PV interneurons from cuprizone-treated mice after bath application of AUT00201 when averaging all the cells together (*group x current two-way repeated measures: n = 13 cells from 4 mice per group: F(13,78)= 0.9280, p = 0.5287. AUT00201 effect: F(1,6)= 5.05, p = 0.0657*).

**Supp. Figure 3.**
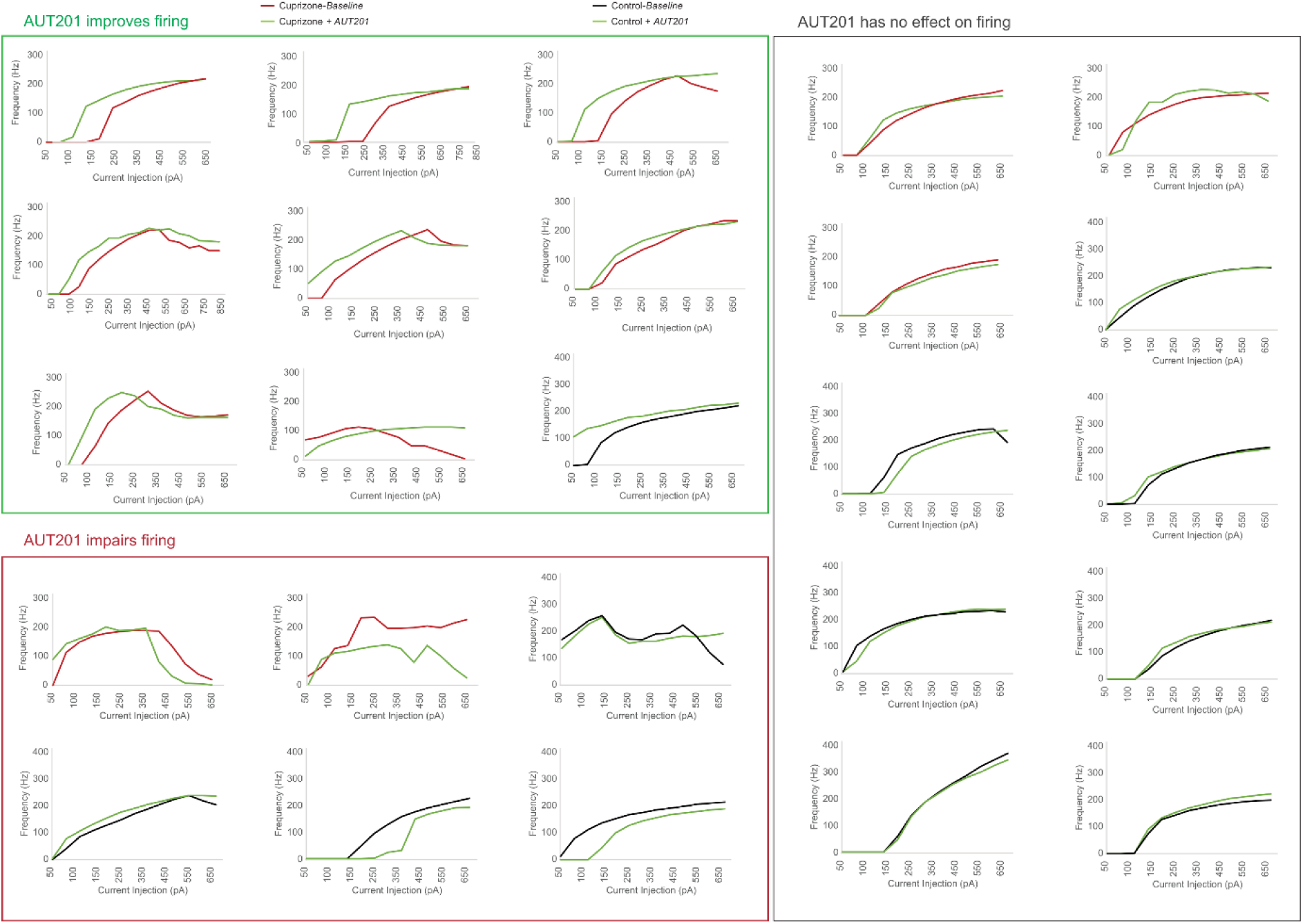
Effect of KV3 modulation on PV interneuron firing frequency and on autaptic release. Effect of bath application of AUT00201 (1uM) (green) on the firing frequency of individual PV interneurons in control (black) and cuprizone-treated (red) mice. The cells were divided into 3 groups: AUT00201 improved the firing, impaired the firing or had no effect.

**Supp. Figure 4.**
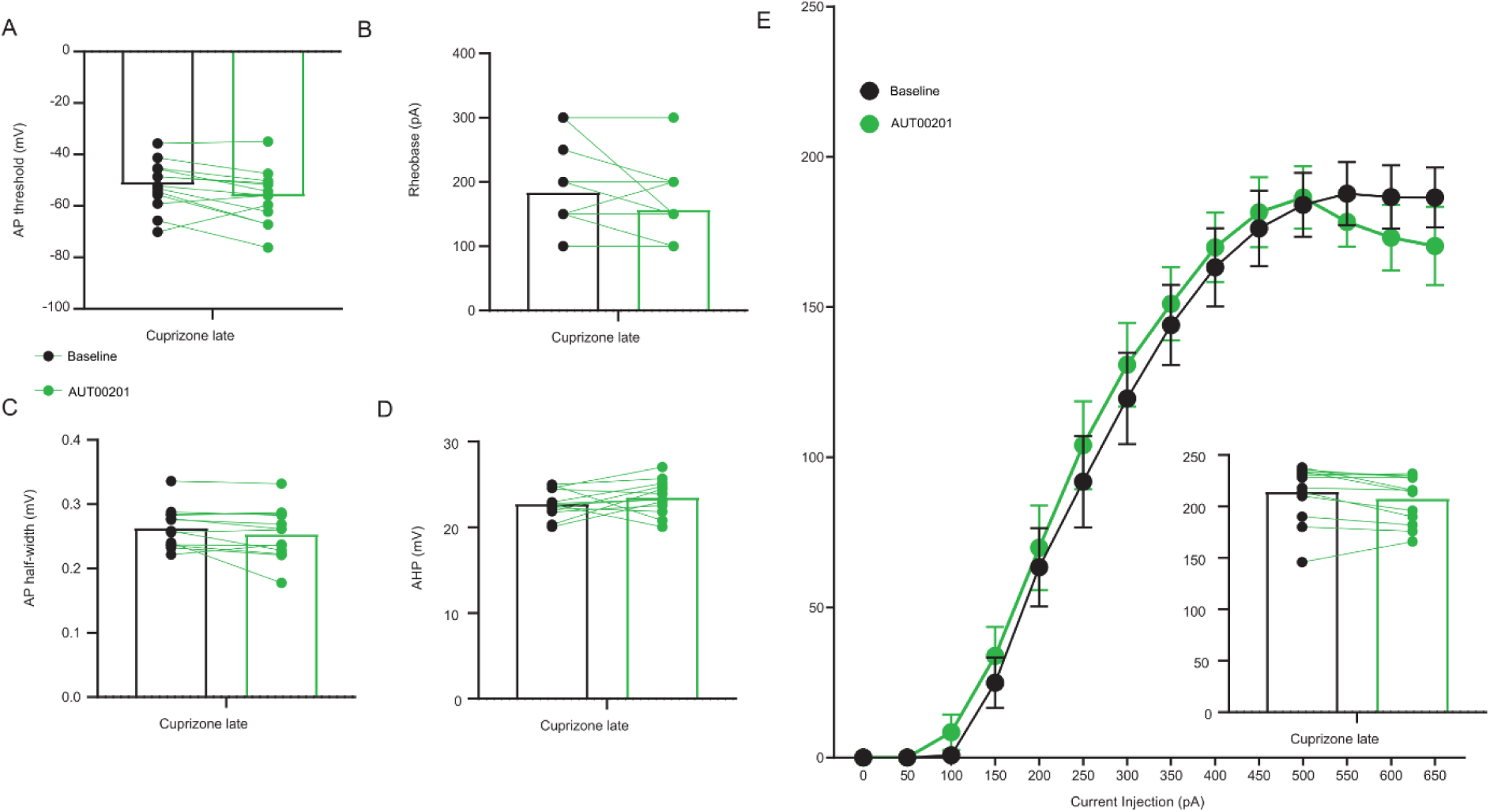
Effect of AUT00201 (1uM) on PV properties after adult demyelination. A-E. Bath application of AUT00201 (1uM) had no significant effect on any of the intrinsic properties of PV interneurons following adult demyelination. Summary data showing the averaged (± s.e.m) of the AP threshold (A) , the rheobase (B) , the AP half-width (C), the AHP (D) and the overall firing frequency of PV interneurons from mice that underwent adult demyelination (E) (*Paired t-test:* A*: p = 0.1006,* B*: p = 0.0678,* C*: p = 0.1006,* D*: p = 0.2812, E: p = 0.0653; two-way repeated measures: AUT00201 effect: F(1,12)= 0.4990, p = 0.4934*).

**Supplementary Figure 5.**
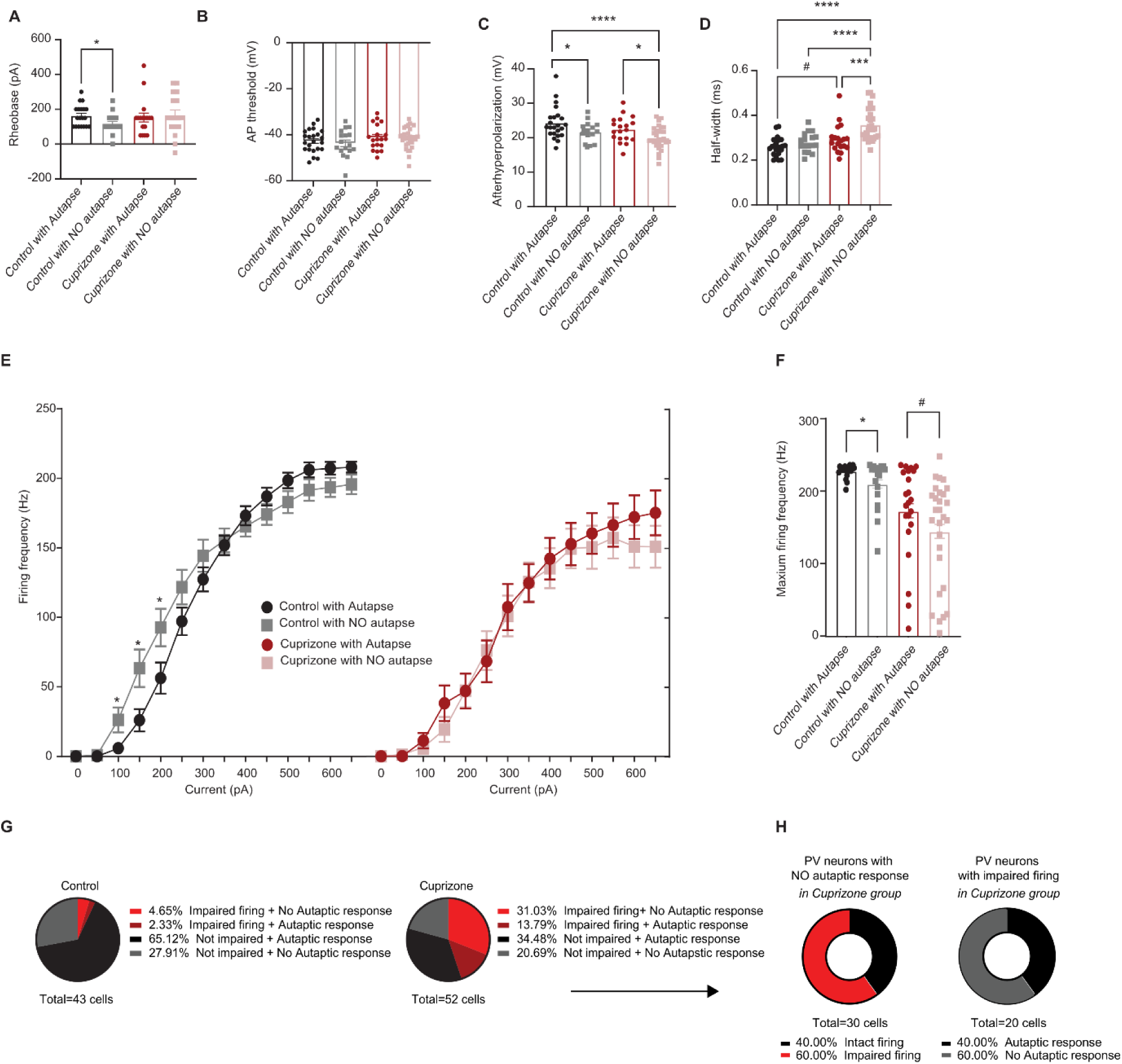
Autapses play a crucial role in PV interneuron’s sustained firing at high frequencies. **A**-**F**. Summary data of the averaged (± s.e.m) altered intrinsic properties of PV interneurons grouped by whether they show autaptic responses or not in both the control and the cuprizone-treated group: (**A**) Rheobase (*t-test of control with vs without autapse; n = 19 cells from 9 mice per group, *p < 0.01; t-test of cuprizone with vs without autapse; n = 17 cells from 8 mice per group, p = 0.6246*), (**B**) Action potential (AP) threshold (*t-test of control with vs without autapse; n = 19 cells from 9 mice per group, p = 0.5680; t-test of cuprizone with vs without autapse; n = 17 cells from 8 mice per group, p = 0.7648*), (**C**) After-hyperpolarization (AHP) amplitude (*ANOVA: n =19,19,18,18 cells from 9-8 mice per group, p = 0.0010, post-hoc LSD test: *p < 0.05, ****p < 0.0001*) and (**D**) AP half-width (*ANOVA: n =19,19,18,18 cells from 9-8 mice per group, p = 0.0002, post-hoc LSD test: #p = 0.0607 ***p < 0.001, ****p < 0.0001*). **E**. Average action potential (AP) frequency in response to 0-650 pA current steps illustrating a significant increase in PV interneuron firing frequency at low-current steps in control cells without autapses (grey) compared to control cells with autapses (black). Note that in cuprizone mice there was no difference (*group x current two-way repeated measures: n = 19,19,18,18 cells from 9-8 mice per group: F(39,856)= 1.671, p = 0.0068, post-hoc LSD test: *p < 0.05*). **F**. Summary data of the maximum firing frequency per group revealing a decrease in the maximum firing frequency in PV interneurons that do not have autapses (*t-test of control with vs without autapse; n = 19 cells from 9 mice per group, *p < 0.01; t-test of cuprizone with vs without autapse; n = 17 cells from 8 mice per group, p = 0.0722*). **G**. Pie chart showing the fraction of cells with impaired or intact firing and with or without autapses in both control mice (left) or cuprizone-treated mice (right). **H.** *Left.* Fraction of cells with intact or impaired firing within all the cells that had no autaptic response in the cuprizone group. *Right.* Fraction of cells with or without an autaptic response within all the cells that had impaired firing in the cuprizone group.

## Notes

### Competing Interest Statement

The authors have declared no competing interest.

### Summary of Updates

The paper was revised after comments from colleagues.

